# SAPID: a Strategy to Analyze Plant Extracts Taste In Depth. Application to the complex taste of *Swertia chirayita* (Roxb.) H.Karst.

**DOI:** 10.1101/2025.01.11.628830

**Authors:** Adriano Rutz, Pascale Deneulin, Ivano Tonutti, Benoît Bach, Jean-Luc Wolfender

## Abstract

Bitterness is challenging to analyze due to the diversity of bitter compounds, variability in sensory perception, and its interplay with other tastes. To address this, we developed an untargeted approach to deconvolute the taste and molecular composition of complex plant extracts. We applied our methodology to an ethanolic extract of *Swertia chirayita* (Roxb.) H.Karst., a plant known for its unique bitterness. Chemical characterization was performed through nuclear magnetic resonance spectroscopy experiments together with untargeted liquid chromatography-high resolution tandem mass spectrometry analysis coupled to a charged aerosol detector. After clustering the fractions based on chemical similarity, we performed free sensory analysis and classical descriptive analysis on each cluster. Our results confirmed the attribution of bitterness to iridoids and highlighted the role of other important compounds in the overall taste. This method offers a systematic approach to analyzing and enhancing the taste profiles of plant-based beverages.

**Highlights:**

- An untargeted method to analyze in depth plant extracts’ taste and chemical composition has been developed.
- The attribution of the bitterness of an ethanolic extract of *Swertia chirayita* (Roxb.) H.Karst. to well-known bitter major and minor iridoids was confirmed using untargeted methods.
- *Chemically informed tasting* allowed to highlight other less pronounced tastes within the extract, contributing to its overall complexity.
- *Chemically informed tasting* led to interesting insights into the sub-threshold impact on taste and taste modulating properties.

**Graphical Abstract:** 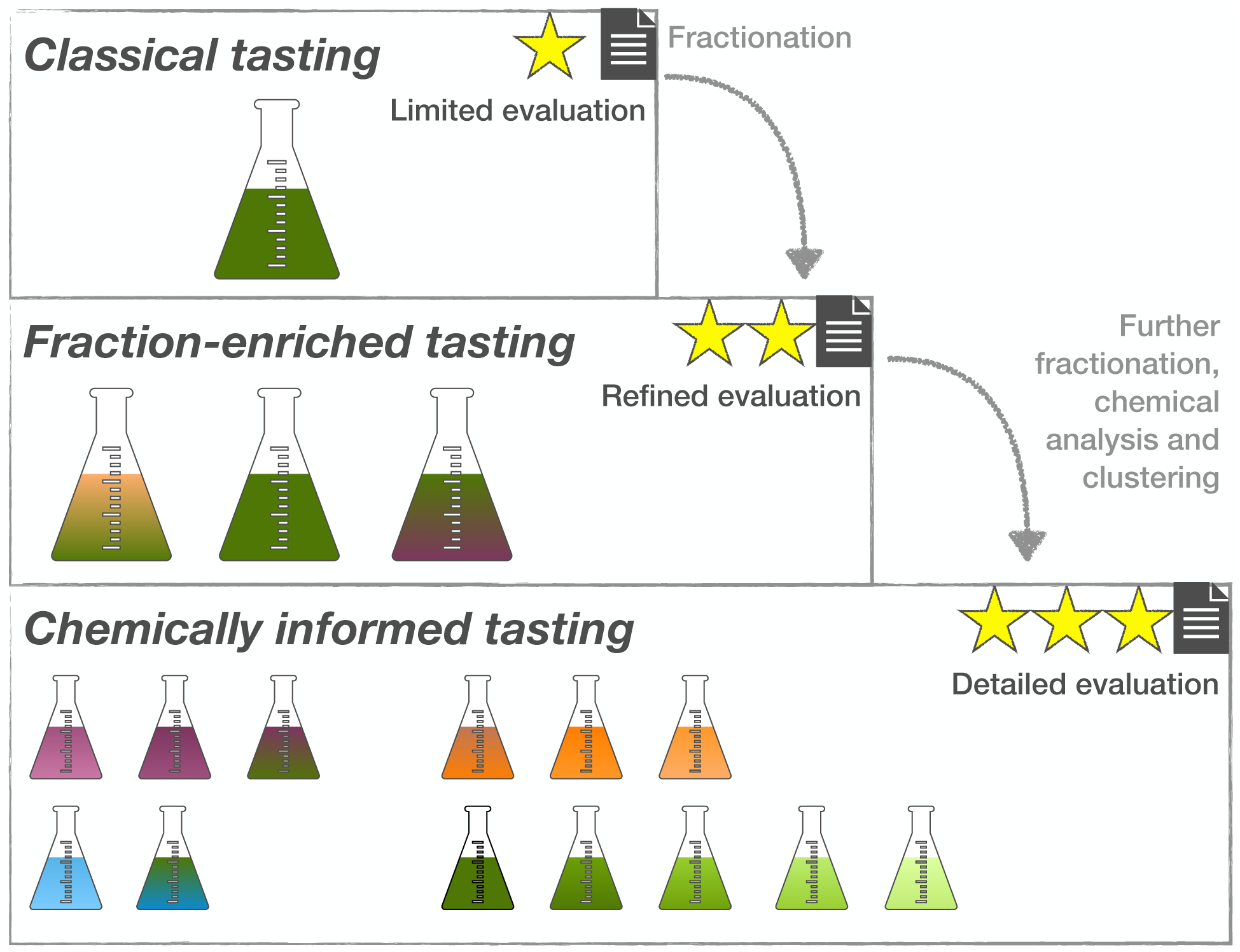

## 1. Introduction

The exploration of bitter compounds in beverages represents a rapidly evolving area of research with significant implications for both the food and beverage industries and human health Joshi, Hankey and Patwardhan (2006). Bitter compounds, found naturally in ingredients such as citrus peels, hops, and various botanicals, contribute to the complexity of flavor profiles that consumers increasingly appreciate in products ranging from craft beers to functional drinks Yan and Tong (2022). While bitterness has historically been avoided in many formulations due to consumer aversion, recent trends show a shift toward a broader acceptance and even preference for bitter flavors, especially within niche markets and health-conscious demographics Belitz and Wieser (1985); Zhao, Jeffery and Miller (2022). This shift has been partly driven by the belief that certain bitter compounds, might offer health benefits Barratt-Fornell and Drewnowski (2002). However, the management of bitterness remains a challenge, as excessive bitterness can lead to product rejection Gaudette and Pickering (2013). Therefore, understanding the sources, perception, and modulation of bitterness in beverages is critical for optimizing sensory appeal while retaining the functional benefits of these bioactive compounds. In this respect, the development of approaches to decipher the taste of natural ingredients at the level of their individual constituents is invaluable in understanding how bitterness is modulated and in helping to optimize the organoleptic properties of final products.

*Swertia chirayita* (Roxb.) H.Karst. is a bitter plant of important medical use in India. It grows between 1200 and 3200m in the Himalayan range Shukla, Bafna, Sundar and Thorat (2014). As it is attributed a significant amount of medicinal properties Saha and Das (2010); Phoboo, Pinto, Barbosa, Sarkar, Bhowmik, Jha and Shetty (2012); Suryawanshi, Mehrotra, Asthana and Gupta (2006) in addition to its organoleptic properties, is a critically endangered species Kumar and Van Staden (2016). Like other Gentianaceae, it is known to contain high amounts of xanthones and iridoids Hostettmann-Kaldas, Hostettmann and Sticher (1981). Among the known iridoids of *S. chirayita* Sharma, Bhardwaj, Sharma, Thakur and Sharma (2022), Amarogentin is one of the most bitter compounds known to date, with a bitter value of 58,000,000 Wölfle and Schempp (2018). The different concentrations and the chemical and sensorial diversity of the compounds found in *S. chirayita*’s extract represent a true analytical challenge.

For these reasons, we decided to further study an enriched ethanolic extract of *S. chirayita* already analyzed in Rutz and Wolfender (2023), to unravel its complex taste, using a *chemically informed tasting* approach. The presented workflow combines fractions enrichment, generic quantitative chemical evaluation through Nuclear Magnetic Resonance (NMR) Spectroscopy, *automated composition assessment* through high resolution tandem mass spectrometry (HRMS/MS) coupled to charged aerosol detection (CAD) and a novel combination of sensory evaluation methods. For the organoleptic aspects, napping Pagès (2003) was used in parallel to free descriptive profiles, and the intensity of the obtained descriptors was also assessed. This untargeted methodology on both sensory and chemical sides should facilitate the deconvolution of extracts with a complex taste.

## 2. Results and Discussion

### 2.1. Fractionation and Clustering

The first necessary step to disentangle the taste of *S. chirayita* in the finest details was to fractionate the extract. Briefly, the raw extract was freeze-dried, and underwent a first fractionation using Vacuum Liquid Chromatography (VLC), as described in 4.3.1. The ethanolic VLC fraction was further fractionated according to the procedure described in 4.3.2 and led to 82 fractions. The recovery was 78% of the initial mass (10 g, table available in Appendix A). 53 out of those 82 fractions were selected for the rest of the study (fractions 17 to 71, the previous ones were empty because of the VLC enrichment procedure used, fraction 49 was empty and fractions above 71 were too apolar). The fractionation allowed at the same time to concentrate the molecules (and thus the tastes) contained in the extract while separating them. This was made to highlight the smallest possible differences, as the tasting sessions were organized so as to taste similar fractions during the same session. Fractions were then analyzed by HRMS/MS and NMR, to gain chemical insights before sensory evaluation to allow *chemically informed tasting*. The similarity of the fractions was evaluated on ^1^H signals, as it offers a generic quantitative chemical view of the major compounds. The detailed clustering of NMR signals is available in Appendix B. The raw extract and obtained fractions were also analyzed by ultra high pressure liquid chromatography (UHPLC)-HRMS/MS-Photo Diode Array(PDA)-CAD. The chromatogram of the enriched extract compared to the chromatograms of the selected fractions is available in Appendix C.

### 2.2. Tasting

To avoid saturation, the panelists could not taste more than 7 fractions per session, organized as follows: First, panelists were given blindly a glass of Chasselas (white wine, not aromatic), the same for each tasting session. They had to score its intensity according to the nine sensory descriptors they were trained on. This had the double advantage of mouth preparation and having a standard for each experiment (a summary of the variation in the scores is available in Appendix D). Then, a group of fractions was evaluated using the ‘napping’ method Pagès (2005). The ‘napping’ method consisted in positioning glasses on a rectangular paper, rectangular paper based on their global similarities or dissimilarities, two glasses are closed to each other if they are perceived as similar. This had the advantage of capturing holistic sensory information that might not otherwise be described. If the panelists could associate a word to describe the position or group of products, they did so and generated a free profile of the fraction. After grouping and cleaning descriptors generated, the vocabulary was then submitted to the panelists in order to score the intensity of each fraction for each descriptor. This tasting has generated a x,y coordinates matrix (coordinates of each sample), eventually added by a scores descriptors matrix, which is compatible with further multivariate data analysis. An example of the whole process and associated statistical treatment is available in Appendix E. Finally, a second glass of the same Chasselas wine was given at the end of the experiment, to assess the possible taste-modulating activity of the fractions. The summary of the taste-modulating activities of each group of fractions is available in Appendix F. In total, among all experiments, over 200 descriptors were generated and curated to be categorized into 14 categories. These descriptors, with their English-translated category, are illustrated in Figure 1.

**Figure 1:**
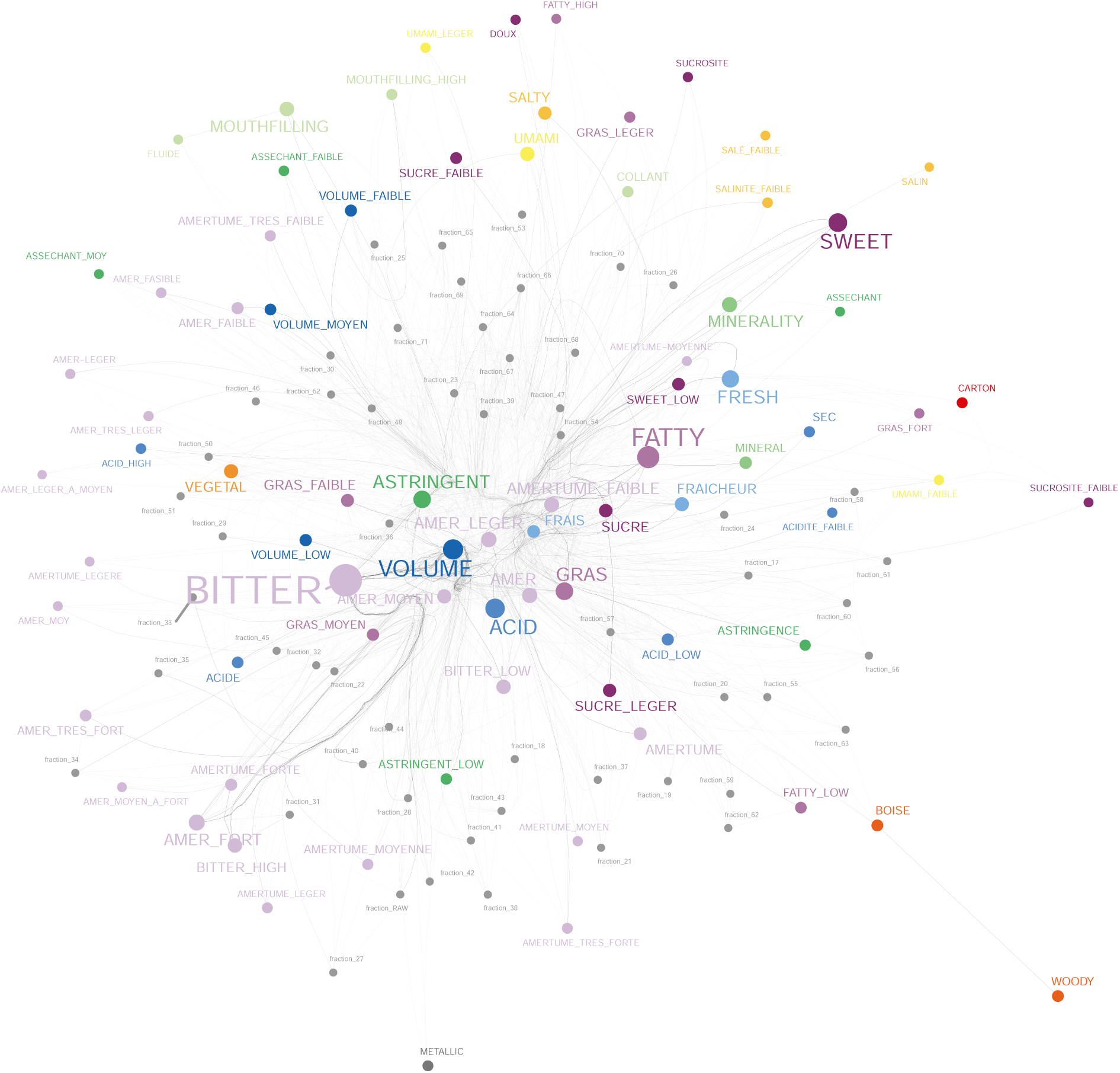
Network of the generated taste descriptors. If two descriptors were used for the same fraction, they are linked together. The color of the edges represents the category to which the descriptor has been assigned. The size of the font and edges represents the number of occurrences. As expected, the category with the highest representation is ‘Bitter’.

The categories illustrated in Figure 1 were used to regroup the different descriptors generated for each group of fractions to gain statistical power. The result of such grouping on the missing values for the bitter taste is available in Appendix G. With those harmonized categories, the intensities reported by each panelist could be summed, to obtain a global view of the taste of each sample.

### 2.3. Sensorial and Chemical Information Gain Due to Fractionation

The result of the classical tasting of the extract using free vocabulary, compared to its *chemically informed tasting* is illustrated in Figure 2.

**Figure 2:**
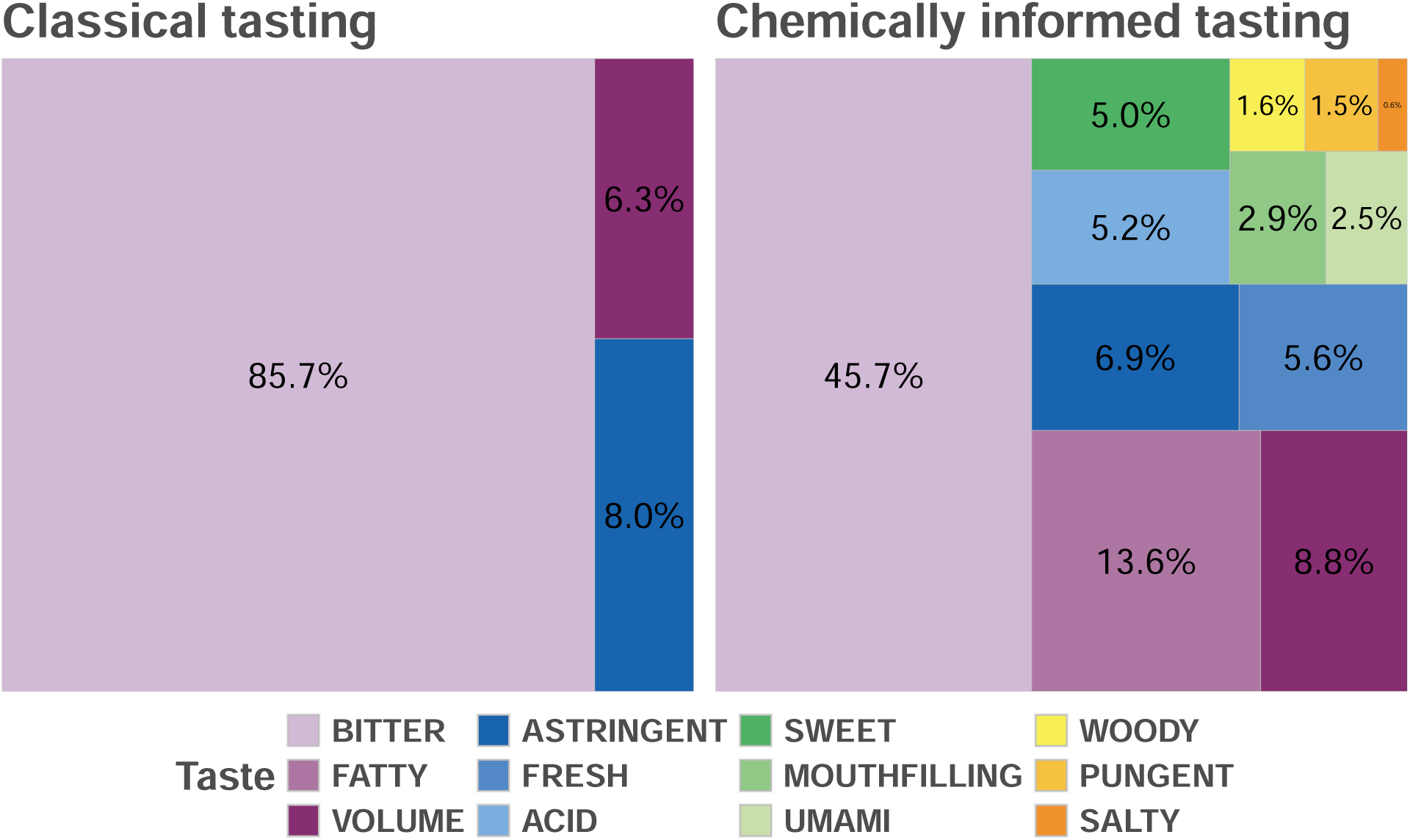
Classical and *chemically informed tasting* of the extract and related fractions. The values represent the sum of the intensities described by at least two panelists, corrected by the mass of the collected fractions. On the left, only three tastes were described for the raw extract, with bitterness accounting for over 85% of the taste intensity. On the right, 9 additional tastes were described when using *chemically informed tasting*, highlighting a richer variety of tastes, contributing to the overall complexity of the extract.

The view presented in Figure 2 illustrates the summary of the information gain about tastes obtained thanks to fractionation, compared to the initial extract. The major taste remained bitterness, but its contribution to the overall taste intensity was almost divided by two (86% to 46%), accounting for the relative masses of the fractions (see Appendix A). While some of the additional tastes detected might seem almost insignificant, the fact that they were reported by two panelists at least consistently with free vocabulary methods clearly shows their importance.

On the chemical side, an *automated composition assessment* was performed on the extract and the fractions Rutz and Wolfender (2023). The fractionation allowed to increase the number of mass spectrometric retained features from 3,471 to 8,104. The features number not only increased but also led to more confident unique annotations (1,562 vs 2,327). Among the 2,327 confident unique annotations from the fractions, 172 structures were already reported in Gentianaceae, of which 63 not originally annotated in the extract. These compounds already found in related organisms were probably not reported yet in *Swertia* spp. because of their low relative amount but could nevertheless play a role in the taste of the extract.

To fully leverage the (un-)annotated ions and their relations to taste(s), their relative amounts along the fractions was further studied, as illustrated in Figure 3. Figure 3 shows the annotated chemical classes and tastes of the fractions. ‘Bitter’ and ‘fatty’ associated categories (see Figure 1) were attributed to all fractions, with bitterness being the most intense, notable in fractions 27-29, 31-36, and 40-43. On the other hand, it is interesting to note that in the fractions between 46 and 54, the bitter taste was clearly less perceived, as shown by the absence of monoterpenoids, and other trigeminal (fresh, astringent, volume) and gustatory (sweet) perceptions are expressed. This chromatographic zone is mainly correlated with the presence of xanthones and their dimers specific to the Gentianaceae family. In the part of the chromatogram where more lipophilic molecules eluted (64-71) mostly triterpenes and fatty acids were annotated, and no specific trend in taste perception was recorded. This probably indicates that such compounds have no effect or a negligible effect on overall taste. Given the diversity of descriptors used, this could also indicate unusual tastes, not consistently recognized among panelists. In fraction 17-26, where the most polar constituents were eluted, no significant bitter taste was perceived although monoterpenoids were present in large amounts (especially 17-21).

**Figure 3:**
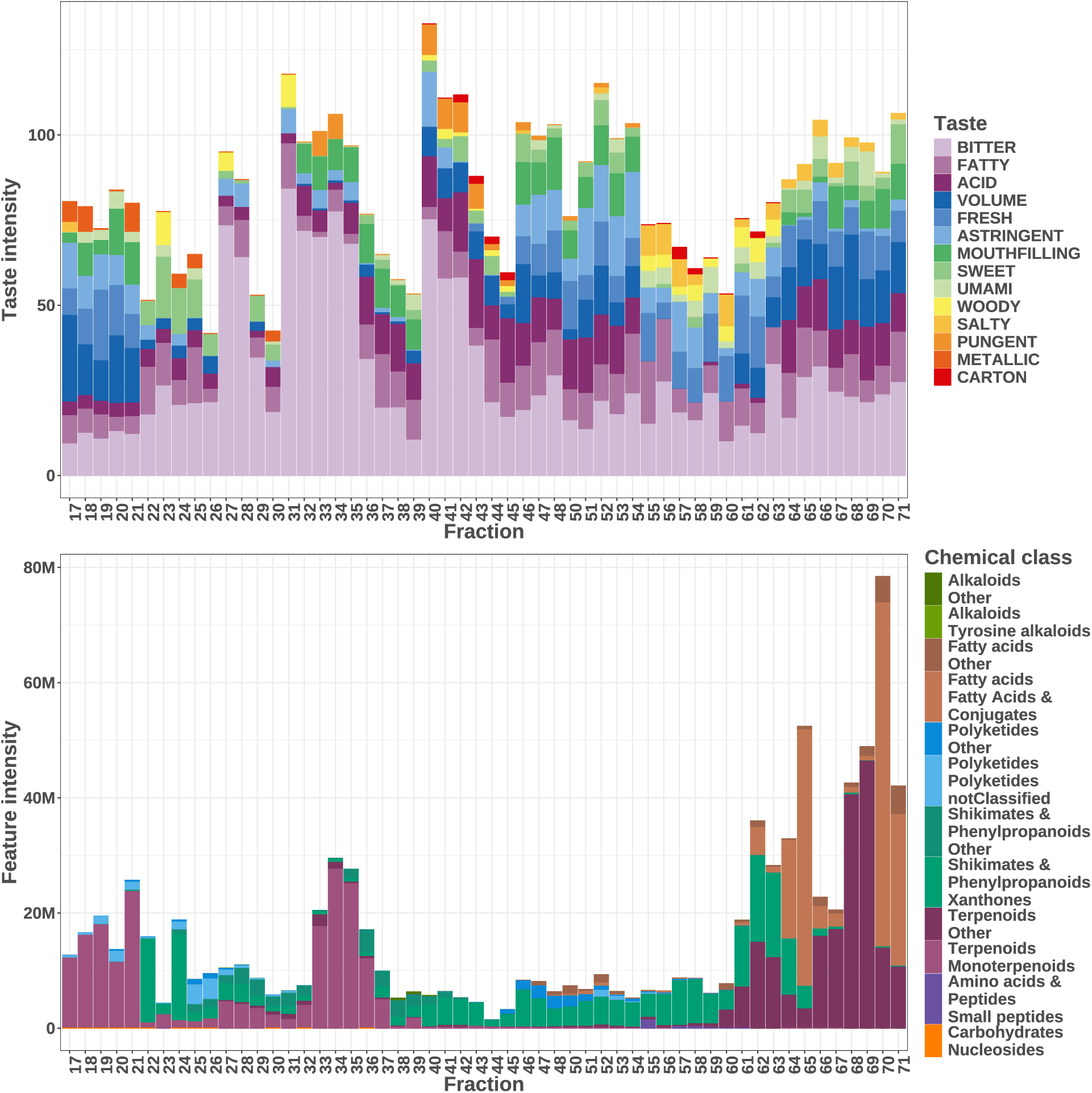
Comparison of the *automated composition assessment* with the *chemically informed tasting* of the fractions. The attributed tastes and their intensity per fraction are shown at the top and the chemical classes at the bottom. At the top, the influence of the tasting session is noticeable. At the bottom, peaks of major terpenoids can be observed.

### 2.4. Sensorial and Chemical Correlations

To refine further the initial trends observed in Figure 3, over 6 million correlations between feature-taste intensity pairs were calculated. This allowed to retain 3,272 correlations (0.05%) as potentially interesting (p-value < 0.05 and correlation > 0.95). Some of the best obtained positive correlations for different tastes are illustated in Figure 4.

**Figure 4:**
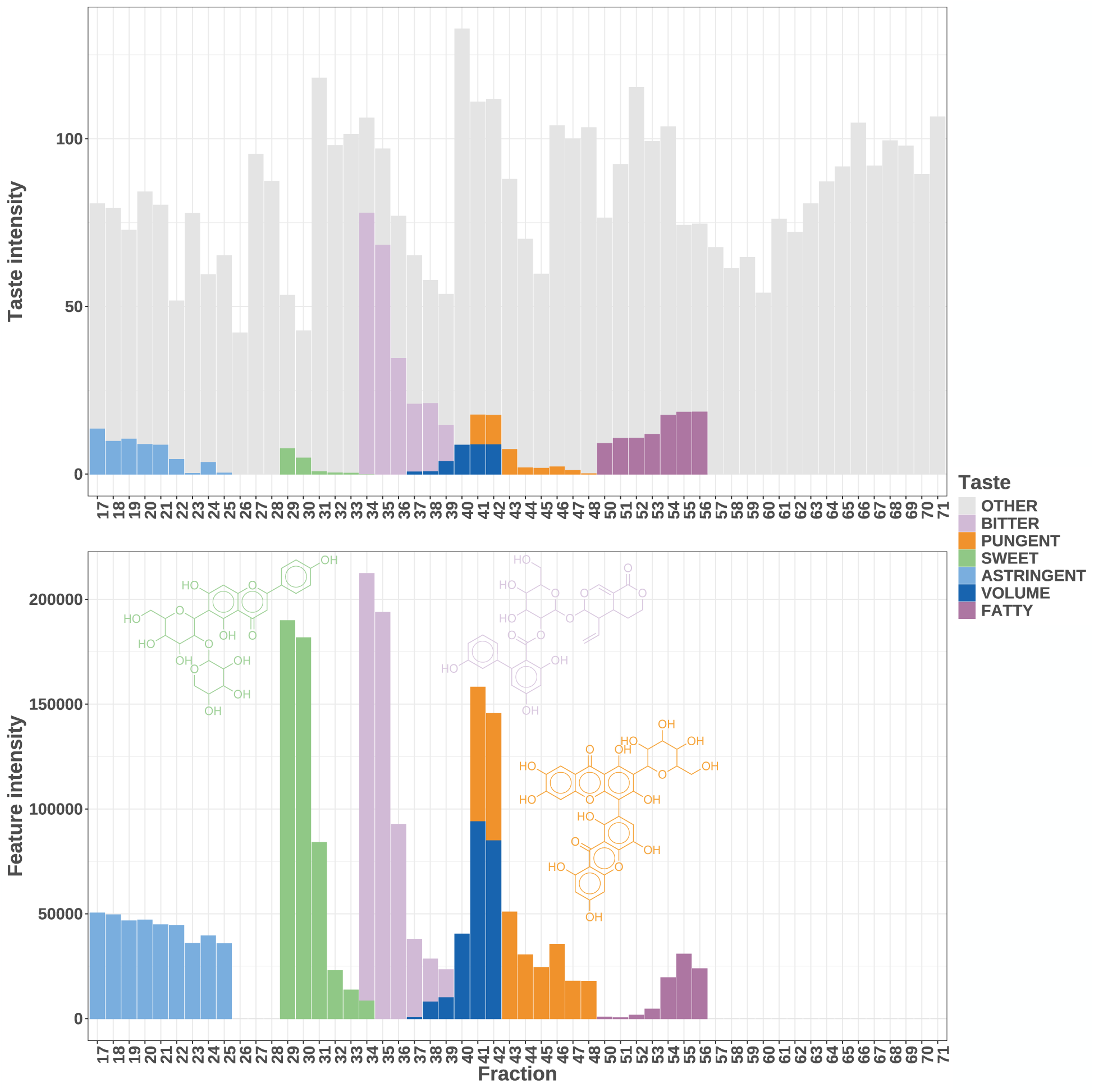
Selected significant feature-taste intensities correlations. The overall strategy allowed, for example, to correlate the bitterness of fractions 34-39 to Amarogentin, the pungency of fractions 41-48 to 3-*O*-Demethylswertipunicoside and sweetness of fractions 29-34 to Isovitexin2”-*O*-arabinoside. Selected correlations are also shown for astringency, volume and fatness but without confident molecular annotation. Correlations could be obtained for different tastes within the same fractions (34: bitter and sweet; 37-39: bitter and volume; 41 and 42: volume and pungent), and among different tasting sessions (see Appendix B). The intensities of the bitter and pungent ions were divided by 300 and 100 for visualization.

In light blue, dark blue, and dark violet, correlations for astringency, volume, and fatness are shown. In light green, despite the overall bitter taste of the extract, an ion annotated as Isovitexin2”-*O*-arabinoside was well correlated to sweet descriptors. To the best of our knowledge, its potential taste has not been reported yet, but its arabinose moiety tastes sweet. Rojas, Ballabio, Pacheco Sarmiento, Pacheco Jaramillo, Mendoza and García (2022) Despite being in a taste-intense region (see Figure 3), 3-*O*-Demethylswertipunicoside was correlated to pungency, one of the least reported taste categories. Finally, as expected, Amarogentin was well correlated to the bitter descriptors of fractions 34-39. As iridoids are known to contribute to the bitterness of Gentianaceae spp. Wolfender, Hamburger, Hostettmann, Msonthi and Mavi (1993), all ions confidently annotated as such were investigated. The results are presented in Figure S7 in Appendix H. The bitterness of the extract correlated well to Amarogentin and Amaroswerin, two well known iridoids of *S. chirayita*, but also to a Decentapicrin derivative. No report of Decentapicrin derivatives bitterness or occurrence in *Swertia* spp. seems to exist to date, but Decentapicrin A was already found together with Amarogentin and Amaroswerin in *Gentianella nitida* Kawahara, Masuda, Sekita and Satake (2001).

It is important to note that the correlations highlighted in Figure 4 are only a small sub-part of the overall obtained correlations. Nevertheless, they illustrate well the strengths and weaknesses of our approach. If two molecules with similar taste elute similarly in close fractions, distinguishing them will remain difficult, and some correlations might be missed, as highlighted by the non-ideal correlation obtained for Amaroswerin in Figure S7. Same applies for fractions where many taste-active molecules are still mixed together, but our approach was still able to prioritize molecules with distinct tastes within the same fractions (fractions 34, 37-39, 41 and 42, Figure 4). Moreover, correlations could be retrieved across different tasting sessions (see Appendix B), thus showing that despite the expected high inter and intra session variance of free vocabulary experiments, meaningful information could be retrieved.

## 3. Conclusion

To identify the compounds responsible for the taste of an ethanolic extract of *S. chirayita*, analytical and sensory experiments were performed, both in an untargeted manner. The extract was fractionated to get a more detailed view of its composition. By taking advantage of the *automated compositional assessment*, an overview of the chemical composition could be obtained. *Chemically informed tasting* allowed the taste-active molecules to be effectively disentangled and their contribution to the overall taste of the extract to be assessed. This could lead to improvements in fine-tuning their selective extraction or synthesis in their usage as flavoring ingredients.

Different iridoids contributed to the bitter taste of the extract, mainly Amarogentin, Amaroswerin, and a Decentapicrin derivative. In addition to these highly bitter molecules, other minor compounds contributed to the overall complex taste of the extract. Interestingly, some fractions showed taste and trigeminal modulatory activity which may contribute to the overall specific bitter perception of the extract. *S. chirayita* could play a valuable role in the production of high-quality bitter beverages.

Further research, such as taste reconstitution and omission experiments Rotzoll, Dunkel and Hofmann (2006); Engel, Nicklaus, Salles and Le Quéré (2002), should be performed to strengthen the results. In the case of rich matrices, these experiments are challenging, as it is almost impossible to achieve complete depletion and residual complexity has been shown to impact results Choules, Klein, Lankin, McAlpine, Cho, Cheng, Lee, Suh, Jaki, Franzblau and Pauli (2018). However, the methodology we developed helps focus and streamline these experimental efforts. These focused efforts should also contribute to fostering a virtuous cycle, where scarce compounds can be utilized to further train panelists. This, in turn, would enhance sensory evaluations, as traditional training methods remain constrained by the limited availability of representative tastant molecules.

While this work was developed with a focus on taste, we believe that our approach will be even more successful with other type of biological activities that are less prone to variation.

## 4. Experimental

### 4.1. Plant Material

The aerial parts of *S. chirayita* were supplied by Tradall SA (Batch 155174).

### 4.2. Extraction

The plant material was used for other studies, therefore high amounts of extract were needed.

1.1 kg of plant material was homogenized in a grinder and extracted at room temperature with 5 kg H_2_O and 6 kg EtOH for 4 days. The extract was then filtered and stored in inox for decantation for 10 days. Finally, the extract was concentrated under reduced pressure, freeze-dried and stored at -20^◦^C until further use. From 1.1 kg of plant material, 86.3 g of dried extract was obtained.

### 4.3. Fractionation

#### 4.3.1. Vacuum Liquid Chromatography

The first fractionation of the extract was undertaken by Vacuum Liquid Chromatography (VLC). 10 g of extract mixed with 20 g of C_18_ silica were loaded on a chromatographic system made of two layers of silica (50 g and 200 g) separated by sand. This system was first very gently (approximately 2 drops/sec) eluted with 3x500 mL 100% H_2_O and then 3x500 mL 100% EtOH. The aqueous and ethanolic parts were collected separately, with a ‘mix’ part corresponding to the dead volume of the system between both and the system washed with DCM. All VLC fractions were then concentrated under reduced pressure, freeze-dried and stored at -20^◦^C until further use. This procedure was repeated 3 times. From the initial 30 g of extract, a total of 27.4 g were recovered (91.3%). Mass of VLC_1 H_2_O was 12.4 g, VLC_2 (mix) was 4.2 g, VLC_3 (EtOH) was 10.4 g and VLC_4 (wash) 0.3 g.

#### 4.3.2. Medium-Pressure Liquid Chromatography

10 g of VLC_3 were mixed with 20 g of Zeoprep® C_18_ (40–63µm, BGB) and filled in an aluminum cartridge. Fractionation was performed using a Buchi 681 pump equipped with a Knauer UV detector and a 460 × 70 mm i.d. column loaded with Zeoprep® C_18_ as stationary phase (15–25µm, BGB). Fractionation was performed at a flow rate of 20 mL/min using a solvent mixture of 0.1% formic acid in water (eluent A) and 0.1% formic acid in methanol (eluent B) and the following gradient: 0 minute, 5% B; held for 37 minutes at 5% B; increased within 600 minutes to 100% B; washed at 100% B. The column effluent was fractionated each 250 mL into 82 sub-fractions (M01–M82). The sub-fractions were then concentrated under reduced pressure, freeze-dried and stored at -20^◦^C until further use. From the initial 10 g of VLC_3, a total of 7.8 g were recovered (77.8%). More details are available in Appendix A.

### 4.4. Sensory Analysis

#### 4.4.1. Panel

The tasting panel included 11 female and 5 male participants. The panelists were trained in sensory evaluation once a week for 2 to 12 years, depending on the participant, before participating in this study. These panelists were primarily trained on wine but they were used to tasting other products. Sensory sessions were run for 10 weeks (once a week) and the number of panelists varied from 9 to 12 per session. Specific training was done for this study.

#### 4.4.2. Organization

The distribution of the samples was done according to proton NMR clustering of the samples. 7 groups were formed by hierarchical clustering to get a reasonable number of samples per session and number of sessions. A cluster was tasted per week. Additionally to the samples of the clusters, panelists were asked to blindly evaluate a Chasselas wine before and after tasting, in order to assess the taste modulating properties of the cluster.

#### 4.4.3. Sample Preparation

To minimize saturation and own ethanol tasting properties, all samples were dissolved in demineralized water. To avoid solubility issues, all samples were first prepared in a 60% EtOH stock solution at 10 mg/mL and then diluted in water.

#### 4.4.4. Working Concentration

To evaluate appropriate working concentration, a first session was dedicated to VLC_3 tasting (ethanolic fraction). After tasting, the working concentration for all experiments was established at 3 mg/L, multiplied by the mass contribution of the Medium Pressure Liquid Chromatography (MPLC) fraction to the VLC_3 fraction. For more details, see Appendix I. For cluster 3, working concentration was divided by 50 as preliminary chemical analyses revealed that they contained extremely bitter compounds. For cluster 7, working concentration was multiplied by 2, as preliminary chemical analyses revealed that it represented the largest amount of the extract’s mass.

### 4.5. Data Acquisition

#### 4.5.1. Mass Spectrometry

Chromatographic separation was performed on a Waters Acquity UHPLC system interfaced to a Corona^TM^ Veo^TM^ RS Charged Aerosol Detector (CAD) and a Q-Exactive Focus mass spectrometer, using a heated electrospray ionization (HESI-II) source. Thermo Scientific Xcalibur 3.1 software was used for instrument control. The conditions were as follows: column, Waters BEH C_18_ 150 × 2.1 mm, 1.7 µm; mobile phase, (A) water with 0.1% formic acid; (B) acetonitrile with 0.1% formic acid; flow rate, 400 µl·min^-1^; injection volume, 6 µl; gradient, isocratic at 5% B for 0.5 minute linear gradient of 5-100% B over 28 minutes and isocratic at 100% B for 12 minutes. The optimized HESI-II parameters were as follows: source voltage, 3.5 kV (pos); sheath gas flow rate N_2_, 55 units; auxiliary gas flow rate, 15 units; spare gas flow rate, 3.0; capillary temperature, 350.00^◦^C, S-Lens RF Level, 45. The mass analyzer was calibrated using a mixture of caffeine, methionine–arginine–phenylalanine–alanine–acetate (MRFA), sodium dodecyl sulfate, sodium taurocholate, and Ultramark 1621 in an acetonitrile/methanol/water solution containing 1% formic acid by direct injection. The data-dependent MS/MS events were performed on the three most intense ions detected in full scan MS (Top3 experiment). The MS/MS isolation window width was 1 Da, and the stepped normalized collision energy (NCE) was set to 15, 30 and 45 units. In data-dependent MS/MS experiments, full scans were acquired at a resolution of 35,000 FWHM (at *m/z* 200) and MS/MS scans at 17,500 FWHM both with an automatically determined maximum injection time. After being acquired in a MS/MS scan, parent ions were placed in a dynamic exclusion list for 2.0 s. A custom exclusion list was used. An Acquity UHPLC PDA detector was used to acquire UV spectra which were detected from 200 to 500 nm. An analytical split was used with a split ratio of 9:1 (CAD:MS). CAD parameters were: evaporation temperature at 40 ^◦^C, 5 bar N_2_, power function 1. The acquisition parameters were the same as described previously Rutz and Wolfender (2023). Analyses were carried in positive mode only.

#### 4.5.2. Nuclear Magnetic Resonance Spectroscopy

NMR experiments (^1^H, ^13^C, and 2D) were performed using a Bruker® Avance III HD 600 (14,1 Tesla) instrument (Bruker BioSpin GmbH, Rheinstetten,Germany) with trimethylsilane (TMS) as internal standard. The details of the sequences are available in the respective subfolders archived on Zenodo (https://zenodo.org/records/14414272) Rutz, Marcourt and Wolfender (2024).

### 4.6. Data Conversion

#### 4.6.1. Mass Spectrometry

All raw data files were converted to .mzML open format Martens, Chambers, Sturm, Kessner, Levander, Shofstahl, Tang, Römpp, Neumann, Pizarro, Montecchi-Palazzi, Tasman, Coleman, Reisinger, Souda, Hermjakob, Binz and Deutsch (2011) using ThermoRawFileParser v.1.4.5 Hulstaert, Shofstahl, Sachsenberg, Walzer, Barsnes, Martens and Perez-Riverol (2019). The generic command used was:

mono ThermoRawFileParser.exe -d $DIRECTORY --allDetectors --format=2

### 4.7. Data Processing

#### 4.7.1. Mass Spectrometry

##### 4.7.1.1. Features’ Extraction

MS features were extracted and informed using mzmine (4.4.0) Pluskal, Castillo, Villar-Briones and Orešič (2010); Heuckeroth, Damiani, Smirnov, Mokshyna, Brungs, Korf, Smith, Stincone, Dreolin, Nothias, Hyötyläinen, Orešič, Karst, Dorrestein, Petras, Du, van der Hooft, Schmid and Pluskal (2024). For the enriched extract, 3 replicates were measured, while fractions were measured once. First, mass detection was performed with a minimal intensity of 1.0E^4^ for MS^1^ and 0 for MS^2^. Chromatograms were built with minimum 4 consecutive scans above 1.0E^4^ and a minimal absolute height of 5.0E^4^. The *m/z* tolerance was set to 12.0 ppm. Chromatograms were then smoothed using Savitzky Golay algorithm and a window of 5. Features were then resolved using the local minimum feature resolver with a chromatographic threshold of 90%, a minimum absolute height of 1.0E^5^, a minimal ratio of peak top over edge of 1.80, a peak duration range from 0.01 to 1.50 minutes and minimum 5 data points. ^13^C isotope filter was then applied using an intra sample tolerance of 6.0 ppm and 0.04 minute. Monotonic shape was required, with a maximum charge of 2. Further, isotopes were searched using again a 6.0 ppm tolerance and a maximum charge of 2. Features lists were then aligned using a 12.0 ppm tolerance with a weight of 3 times the one of 0.15 minute. Only features with a valid isotopic pattern and at least one associated fragmentation spectrum were retained. For both the 3 extract replicates and the fractions, features had to be present in at least 3 samples. Gap filling was then performed using an intensity tolerance of 20%, 12.0 ppm and 0.15 minute. The list was further refined, removing duplicates using a tolerance of 6.0 ppm and 0.15 minute. For fractions, features present in all samples were removed. Features were then correlated using a 0.08 minute tolerance, an intensity threshold of 1.0E^5^, a minimum of 5 data points, of which 2 on edge. Correlation measure was PEARSON on both shape and height with a minimal correlation of 65%. Ion identity networking Schmid, Petras, Nothias, Wang, Aron, Jagels, Tsugawa, Rainer, Garcia-Aloy, Dührkop, Korf, Pluskal, Kameník, Jarmusch, Caraballo-Rodríguez, Weldon, Nothias-Esposito, Aksenov, Bauermeister, Albarracin Orio, Grundmann, Vargas, Koester, Gauglitz, Gentry, Hövelmann, Kalinina, Pendergraft, Panitchpakdi, Tehan, Le Gouellec, Aleti, Mannochio Russo, Arndt, Hübner, Hayen, Zhi, Raffatellu, Prather, Aluwihare, Böcker, McPhail, Humpf, Karst and Dorrestein (2021) was performed using a maximum charge of 2, a maximum molecules per cluster of 2, and the following adducts list: [M+H]^+^,[M+NH_4_]^+^,[M+Na]^+^,[M-H+2Na]^+^,[M+2H]^2+^,[M+H+NH_4_]^2+^,[M+H+Na]^2+^, together with the following modifications list: [M-C_6_H_10_O_5_],[M-C_6_H_10_O_4_], [M-2H_2_O],[M-H_2_O], [M-NH_3_], [M-CH_3_], [M-C_2_H_3_N]. Finally, ion identities were added using a two step approach, first with an extended range of adducts, then with an extended range of modifications (see batch file). All parameters were given in the form of a batch file, archived within the project’s main repository as described later.

##### 4.7.1.2. Features’ Annotation

Features were annotated using a multi-tool approach already described in Rutz, Dounoue-Kubo, Ollivier, Bisson, Bagheri, Saesong, Ebrahimi, Ingkaninan, Wolfender and Allard (2019); Rutz and Wolfender (2023).

###### 4.7.1.2.1. Sirius

Sirius annotations were performed in batch mode using Sirius 6.0.7 Dührkop, Fleischauer, Ludwig, Aksenov, Melnik, Meusel, Dorrestein, Rousu and Böcker (2019). Compounds above 800 *m/z* were omitted annotation was performed using Orbitrap default parameters. Adducts were the ones already defined in the previous subsection. Formulas were annotated using bottom-up approach, with default parameters Xing, Shen, Xu, Li and Huan (2023). Formulas were annotated using ZODIAC, with default parameters Ludwig, Nothias, Dührkop, Koester, Fleischauer, Hoffmann, Petras, Vargas, Morsy, Aluwihare, Dorrestein and Böcker (2020). CSI:FingerID fingerprints were computed using default parameters Dührkop, Shen, Meusel, Rousu and Böcker (2015). Compound classes were annotated using CANOPUS using default parameters Dührkop, Nothias, Fleischauer, Reher, Ludwig, Hoffmann, Petras, Gerwick, Rousu, Dorrestein and Böcker (2020). Confidence scores were calculated through COSMIC Hoffmann, Nothias, Ludwig, Fleischauer, Gentry, Witting, Dorrestein, Dührkop and Böcker (2021). De novo structure annotation was also performed using MSNovelist Stravs, Dührkop, Böcker and Zamboni (2022).

###### 4.7.1.2.2. In Silico Library

The in silico library used was generated as described in Allard, Bisson and Rutz (2023). The SMILES used were coming from Rutz, Bisson and Allard (2023). CFM 4.0 was used for in silico fragmentation Wang, Liigand, Tian, Arndt, Greiner and Wishart (2021). Parameters are available at https://github.com/mandelbrot-project/spectral_lib_builder.

###### 4.7.1.2.3. Taxonomically Informed Metabolite Annotation

Taxonomically Informed Metabolite Annotation was performed using all the above mentioned inputs. It was performed using TIMA (2.11.0) with default parameters, and a maximum number of candidates of 500 Rutz et al. (2019); Rutz (2024b). The resulting parameters files were archived within the project’s main repository as described later.

##### 4.7.1.3. Features’ Correlation

Because of its sensitivity, and because minor compounds might also have a very strong impact on taste, MS was chosen over NMR or CAD to calculate correlations to the reported taste intensities.

###### 4.7.1.3.1. Correlation to Charged Aerosol Detector

The extracted MS features were used to extract their related MS peak shapes and compare them to the CAD peaks detected in the same retention time window, as described in Rutz and Wolfender (2023). Default parameters from cascade (v.0.0.0.9000) were used. Peaks were detected using routine functions imported from chromatographR package Bass (2023). Peak shapes were then compared using the compareChromatograms function from the MSnbase package with the closest method as argument Gatto, Gibb and Rainer (2020).

###### 4.7.1.3.2. Correlation to Taste

Only tastes reported at least by two panelists were kept, and their intensities summed. Only ions detected in at least 5 fractions were kept. Missing values were imputed using half of the lowest value. Then, MS intensities and taste intensities were correlated using the following steps: First, segments of fractions including 5 to 9 fractions were generated. For each one of these segments, the distribution of taste and ion intensities were evaluated using a Shapiro-Wilk test Royston (1982). If they significantly varied from a normal distribution (p-value < 0.05), correlation was computed using Kendall’s τ, else it was computed using Pearson’s coefficient Hollander, A. Wolfe and Chicken (2015). P-values were then adjusted using Benjamini-Hochberg method Benjamini and Hochberg (1995). The correlations were refined to include only those where the evaluated ion’s intensity ranked among the top five intensity values of the ion at least four times.

#### 4.7.2. Nuclear Magnetic Resonance Spectroscopy

The data were processed using AlpsNMR (4.8.0) Madrid-Gambin, Oller-Moreno, Fernandez, Bartova, Giner, Joyce, Ferraro, Montoliu, Moco and Marco (2020). The ppm values of samples 69, 70 and 71 were manually fixed as they deviated significantly from the rest of the experiment. Then, signals were phased automatically, using background correction and default parameters. Signals were then interpolated from 0 to 14.85 ppm using automatically calculated resolution. Peaks were then detected using lorentzian fits, signals normalized, and peaks integrated using a peak width of 0.01 ppm. Finally, samples were clustered on the square root of their peak areas, using canberra distance measure Lance and Williams (1966) and ward.D2 agglomeration method Ward (1963). 7 clusters were formed from the resulting dendrogram and used further to organize tasting sessions.

### 4.8. Code Availability

All programs written for this work can be found in the following repository: https://github.com/Adafede/sapid. Version 0.0.0.9000 was archived on Zenodo (https://zenodo.org/records/14616396) Rutz (2025).

Main dependencies were AlpsNMR (4.8.0) Madrid-Gambin et al. (2020), cascade (0.0.0.9000) Rutz and Wolfender (2023); Rutz (2024a) (relying heavily on MSnbase Gatto et al. (2020) and Spectra Rainer, Vicini, Salzer, Stanstrup, Badia, Neumann, Stravs, Verri Hernandes, Gatto, Gibb and Witting (2022)) dendextend (1.19.0) Galili (2015), FactoMineR (2.11) Lê, Josse and Husson (2008), forcats (1.0.0) Wickham (2023), ggbump (0.1.0) Sjoberg (2020), ggplot2 (3.5.1) Wickham (2016), ggpubr (0.6.0) Kassambara (2023), ggraph (2.2.1) Pedersen (2024), igraph (2.1.2) Csardi and Nepusz (2006), khroma (1.14.0) Frerebeau (2024), NMRphasing (1.0.5) Jiang (2024), purrr (1.0.2) Vaughan and Dancho (2022), readxl (1.4.3) Wickham and Bryan (2023), scales (1.3.0) Wickham, Pedersen and Seidel (2023), SensoMineR (1.27) Husson, Le and Cadoret (2023), stringi (1.8.4) Gagolewski (2022), tibble (3.2.1) Müller and Wickham (2023), tidytable (0.11.1) Fairbanks (2024), treemapify (2.5.6) Wilkins (2023).

### 4.9. Data Availability

The sensorial data were converted to open format and archived on Zenodo (https://zenodo.org/records/14616396) Rutz (2025).

The raw NMR data were archived on Zenodo (https://zenodo.org/records/14414272) Rutz et al. (2024), as recommended by McAlpine, Chen, Kutateladze, MacMillan, Appendino, Barison, Beniddir, Biavatti, Bluml, Boufridi, Butler, Capon, Choi, Coppage, Crews, Crimmins, Csete, Dewapriya, Egan, Garson, Genta-Jouve, Gerwick, Gross, Harper, Hermanto, Hook, Hunter, Jeannerat, Ji, Johnson, Kingston, Koshino, Lee, Lewin, Li, Linington, Liu, McPhail, Molinski, Moore, Nam, Neupane, Niemitz, Nuzillard, Oberlies, Ocampos, Pan, Quinn, Reddy, Renault, Rivera-Chávez, Robien, Saunders, Schmidt, Seger, Shen, Steinbeck, Stuppner, Sturm, Taglialatela-Scafati, Tantillo, Verpoorte, Wang, Williams, Williams, Wist, Yue, Zhang, Xu, Simmler, Lankin, Bisson and Pauli (2019), .

The raw UHPLC-MS/MS-PDA-CAD data were converted to open format and archived on MassIVE (https://massive.ucsd.edu/ProteoSAFe/dataset.jsp?accession=MSV000096654), together with ReDU compliant metadata Jarmusch, Wang, Aceves, Advani, Aguirre, Aksenov, Aleti, Aron, Bauermeister, Bolleddu, Bouslimani, Caraballo Rodriguez, Chaar, Coras, Elijah, Ernst, Gauglitz, Gentry, Husband, Jarmusch, Jones, Kamenik, Le Gouellec, Lu, McCall, McPhail, Meehan, Melnik, Menezes, Montoya Giraldo, Nguyen, Nothias, Nothias-Esposito, Panitchpakdi, Petras, Quinn, Sikora, van der Hooft, Vargas, Vrbanac, Weldon, Knight, Bandeira and Dorrestein (2020).

## Acknowledgments

The authors gratefully acknowledge l’école du vin in Changins, and (Eve Dante, Amandine Pasquier, and Pierrick Rébénaque-Martinez and all panelists). The authors also thank Magdalena Rausch and Jonathan Bisson for providing constructive feedback. The authors also thank Laurence Marcourt for the NMR data acquisition. J-LW is thankful to the Swiss National Science Foundation for the support in the acquisition of the NMR 600 MHz (SNF R’Equip grant [316030_164095]). This research was funded in whole or in part by the Swiss National Science Foundation (SNSF) [CRSII5_189921]. For the purpose of Open Access, a CC BY public copyright licence is applied to any Author Accepted Manuscript (AAM) version arising from this submission.

## CRediT authorship contribution statement

Adriano Rutz: Conceptualization, Data curation, Formal analysis, Investigation, Methodology, Project administration, Software, Supervision, Validation, Visualization, Writing – original draft, Writing – review and editing . Pascale Deneulin: Methodology, Resources, Supervision, Writing – review and editing . Ivano Tonutti: Funding acquisition, Resources . Benoît Bach: Resources, Writing – review and editing. Jean-Luc Wolfender: Funding acquisition, Resources, Supervision, Writing – review and editing.

## 5. Appendices

### A. Masses of the MPLC fractions

### B. ^1^H NMR of the MPLC fractions

### C. Chromatogram of the enriched extract compared to the chromatograms of the MPLC fractions

### D. Variation of the scoring of Chasselas among all experiments

### E. Initial analysis of the sensory results

### F. Summary of the taste modulating activity for each group of fractions

### G. Matrices of fractions reported as bitter before and after vocabulary curation

### H. Correlations of the intensities of features confidently annotated as iridoids and bitter taste

### I. Determination of the concentration used for tasting

**Figure S1:**
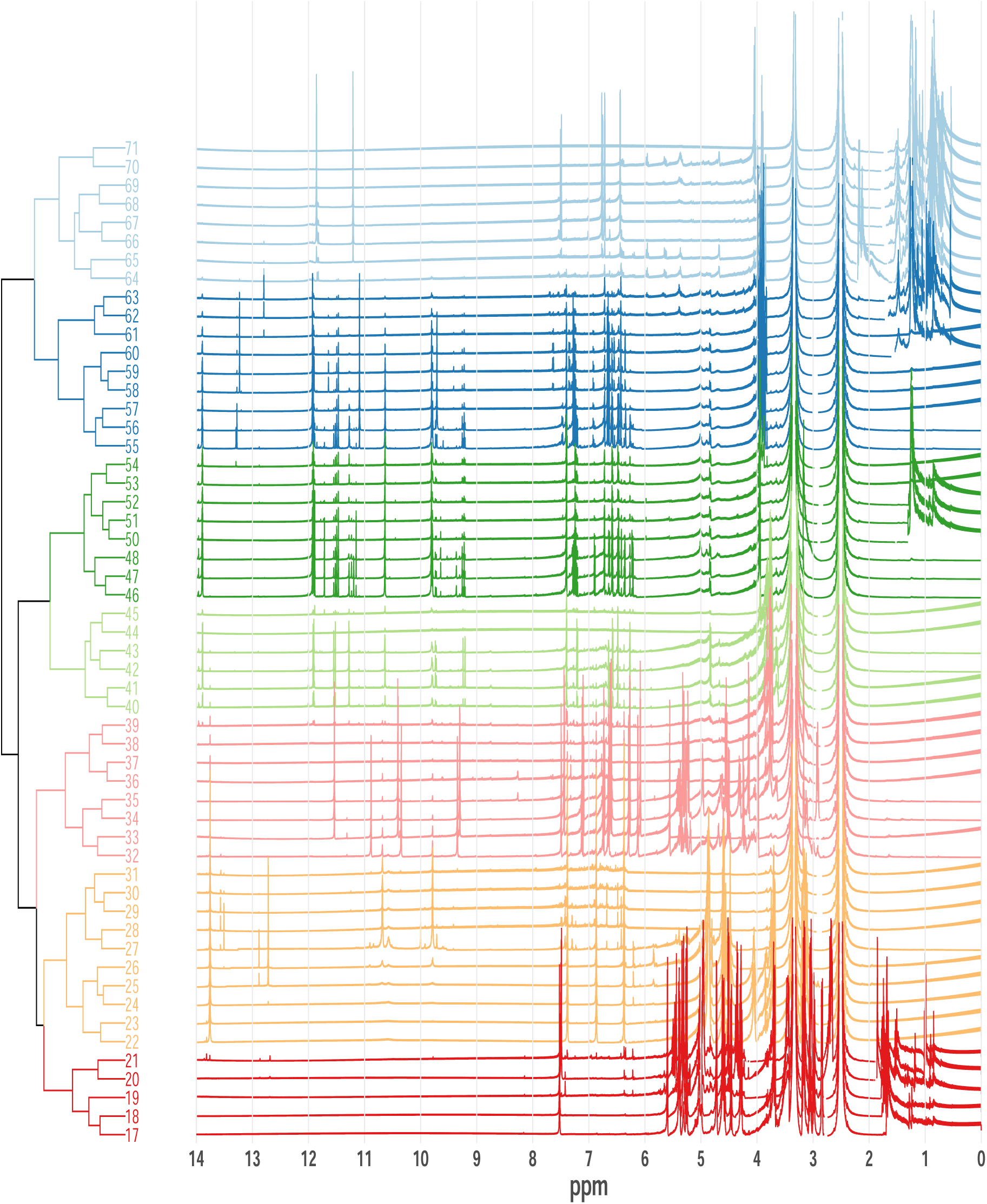
^1^H NMR of the MPLC fractions. Fractions are colored by the group they belong to after clustering. In this case, clusters followed chromatographic order, but this must not necessarily be the case.

**Figure S2:**
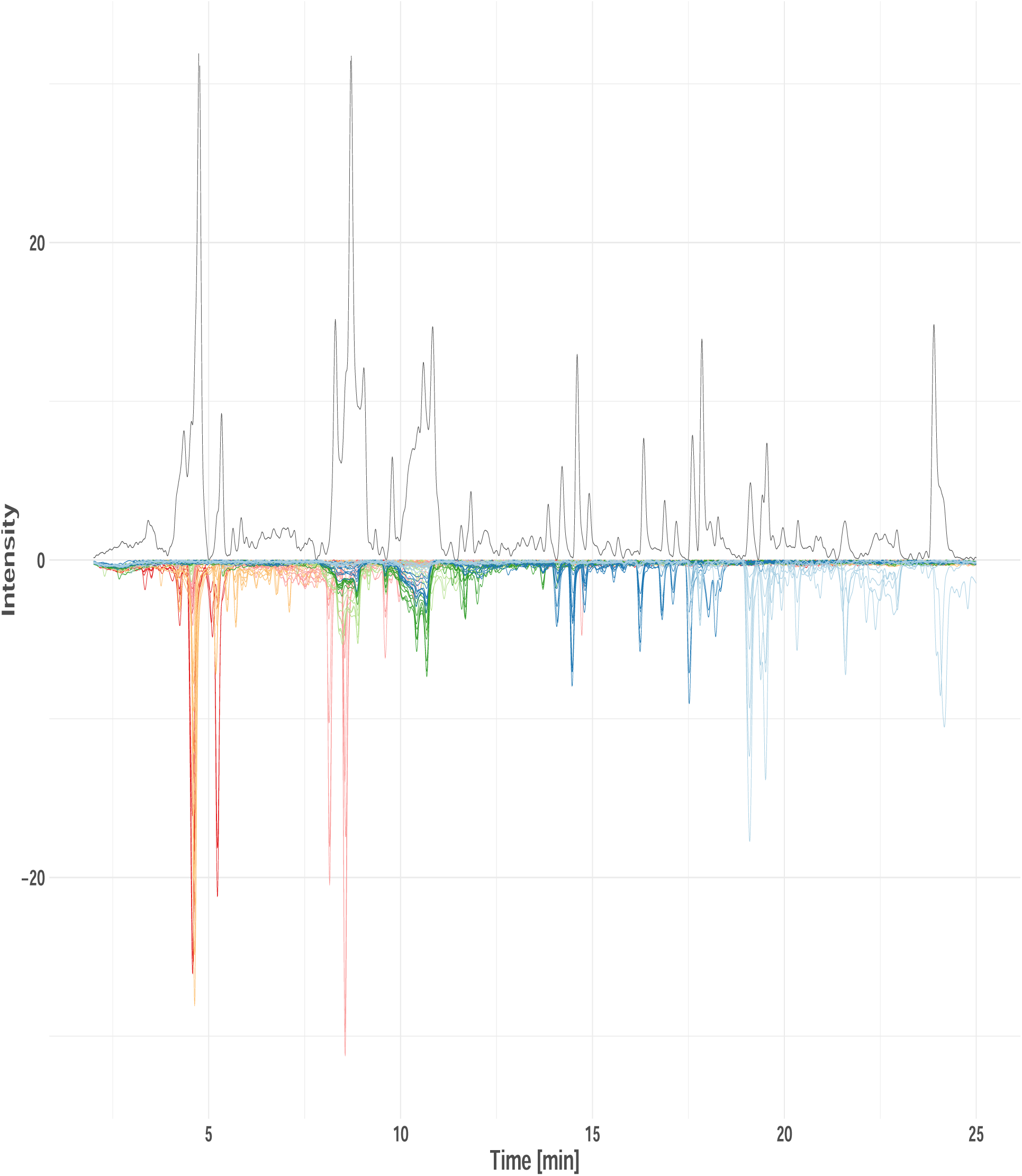
Chromatogram of the enriched extract compared to the chromatograms of the MPLC fractions. The chromatogram of the enriched extract is presented on the top, whereas the chromatograms of the MPLC fractions are presented on the bottom. MPLC fractions are colored according to the clustering made in Appendix B. CAD signal preprocessed as described in Rutz and Wolfender (2023). Except for the peaks at the end of the chromatogram (those fractions were discarded as being too apolar), the profiles of the enriched extract and the fractions look almost identical.

**Figure S3:**
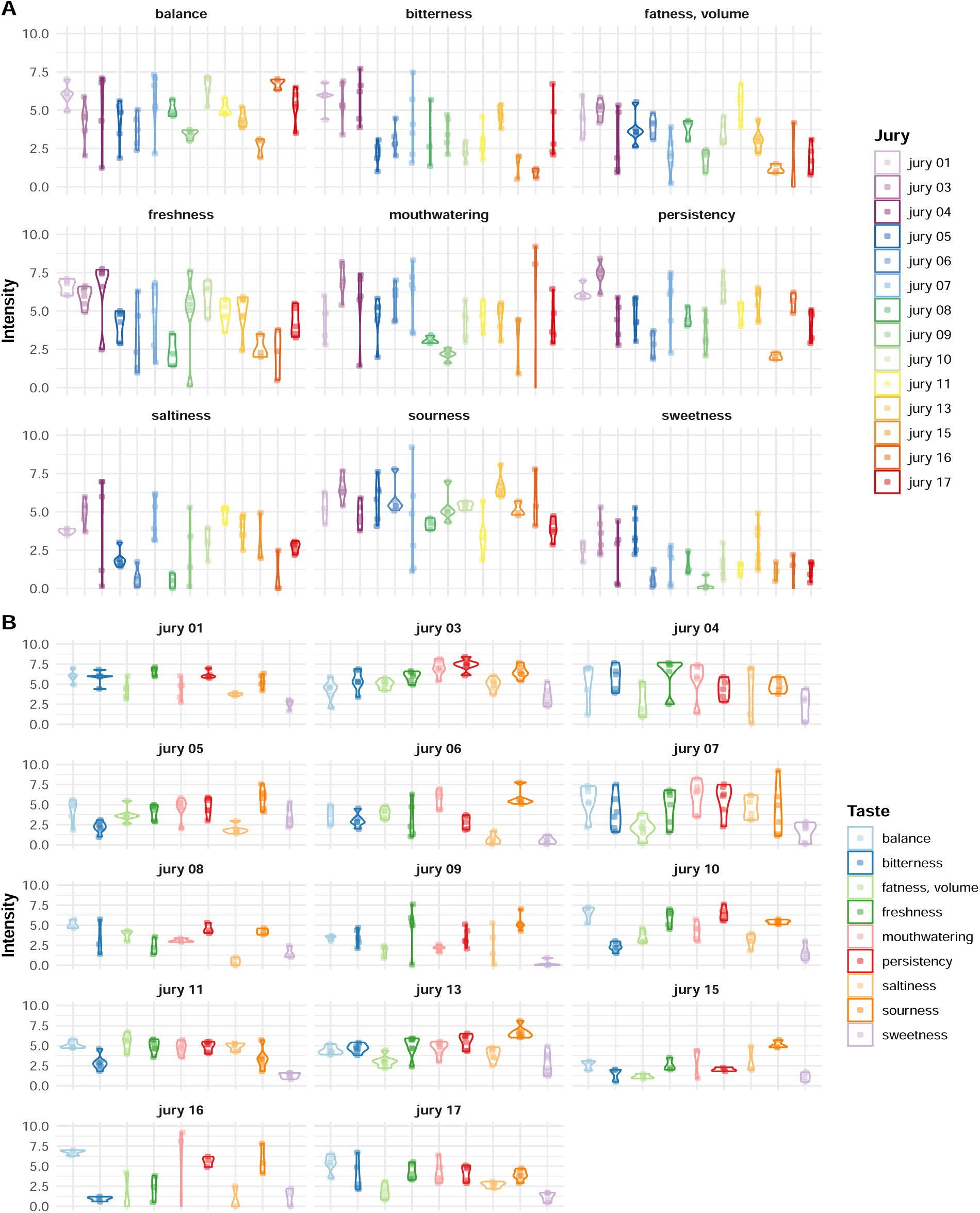
Variation of the scoring of Chasselas among all experiments. In panel A, variations are faceted by taste with jury as color. In panel B, variations are faceted by jury with taste as color.

**Figure S4:**
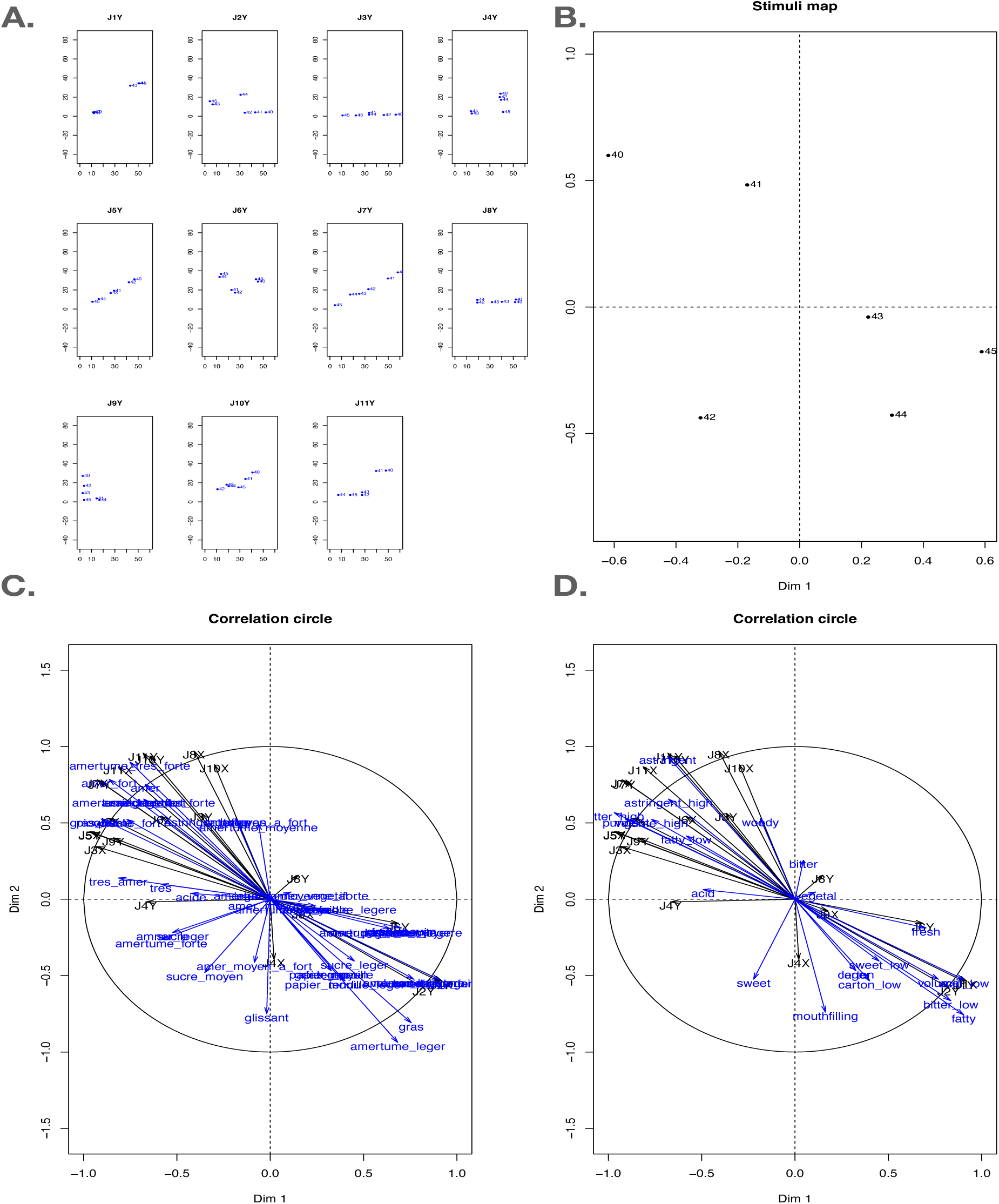
Initial analysis of the sensory results. The positions of each fraction for each panelist are shown in panel A. Using the INDSCAL model Husson and Pagès (2006), a summarized stimuli map is obtained (panel B). This map can then be compared to the correlation circle of the attributed descriptors (panel C). Eventually, a curation of the vocabulary used can occur, as in panel D.

**Figure S5:**
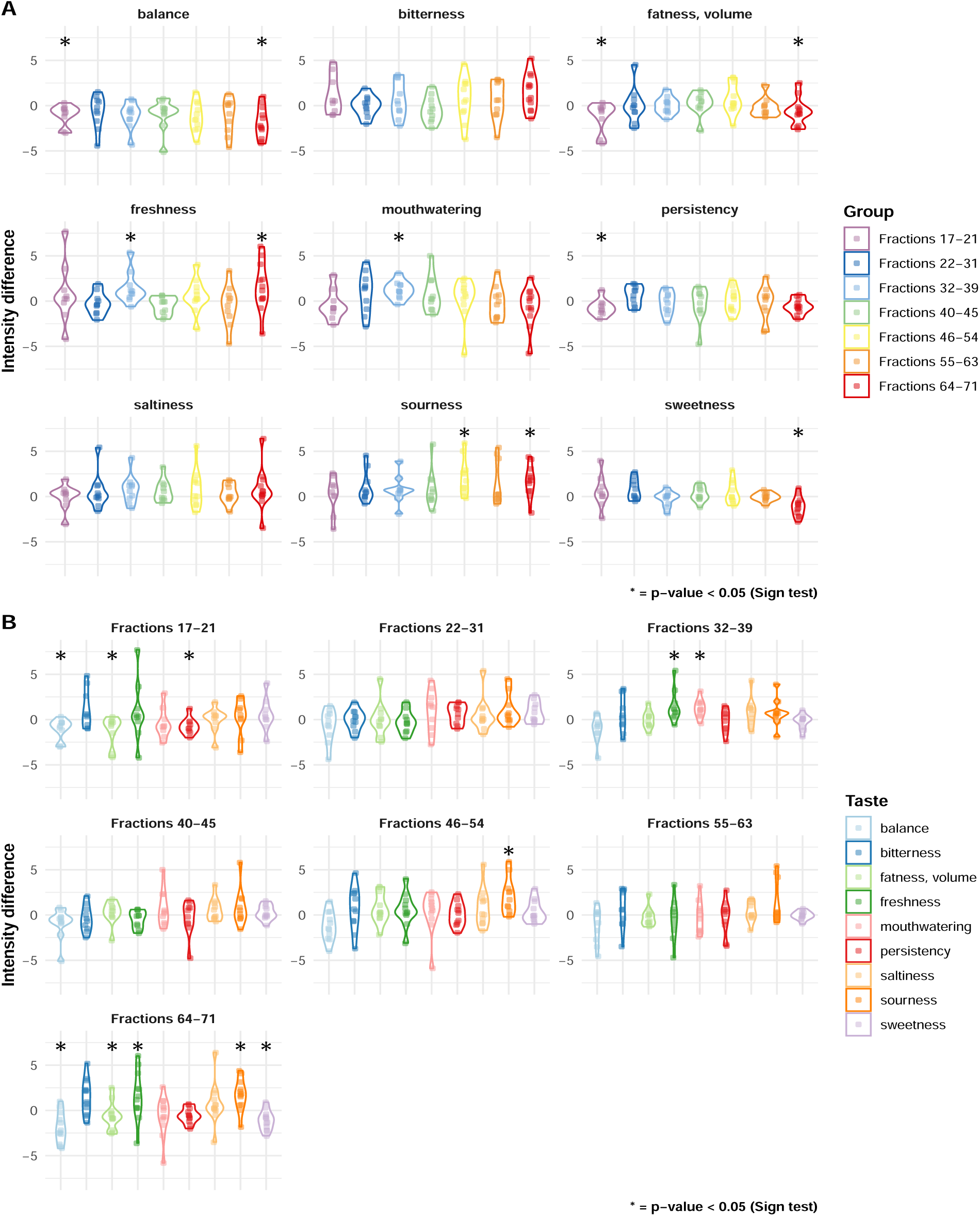
Summary of the taste modulating activity for each group of fractions. The values correspond to the score given to the Chasselas sample tasted after fractions tasting in comparison to the one tasted before fractions. In panel A, variations are grouped by experiment with taste as the variable. In panel B, variations are grouped by taste with experiment as the variable. Stars denote statistical significance according to Sign test (https://en.wikipedia.org/wiki/Sign_test).

**Figure S6:**
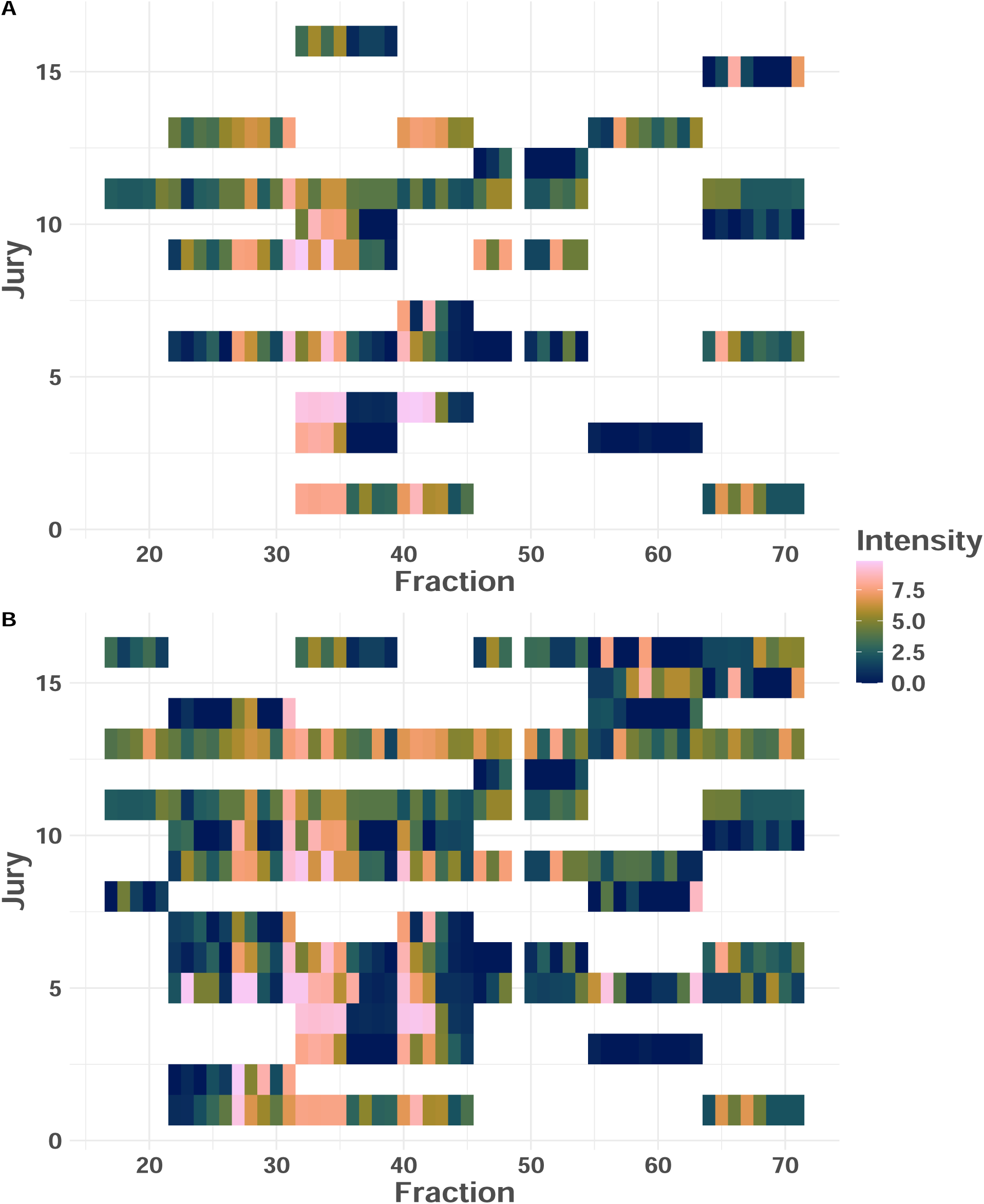
Matrices of fractions reported as bitter before and after vocabulary curation. In panel A, values correspond to the term ‘amer’ (bitter in French) only. In panel B, terms related to bitterness were grouped together as illustrated in Figure 1

**Figure S7:**
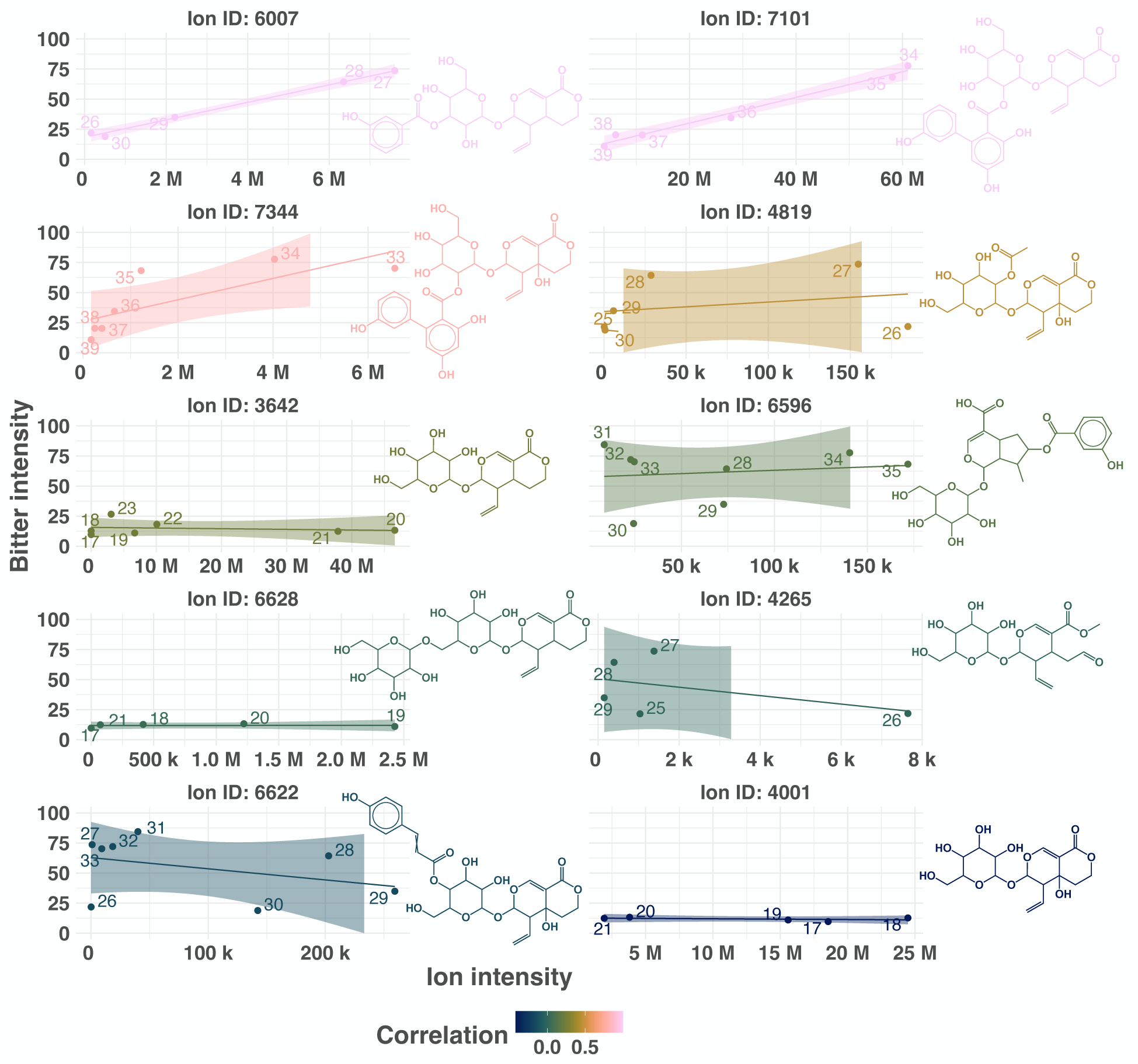
Correlations of the intensities of features confidently annotated as iridoids and bitter taste. The two first ions, annotated as a Decentapicrin derivative (6007) and Amarogentin (7101) respectively are well correlated to bitterness. The ion annotated as Amaroswerin (7344) has a correlation of 0.81, while other ions annotated as iridoids are not correlated, and thus probably not contributing to the bitter taste of the extract. While the presented set of 10 iridoids is not sufficient to draw any conclusions, interesting trends can be observed and larger sets could lead to interesting structure-activity relationships.

**Figure S8:**
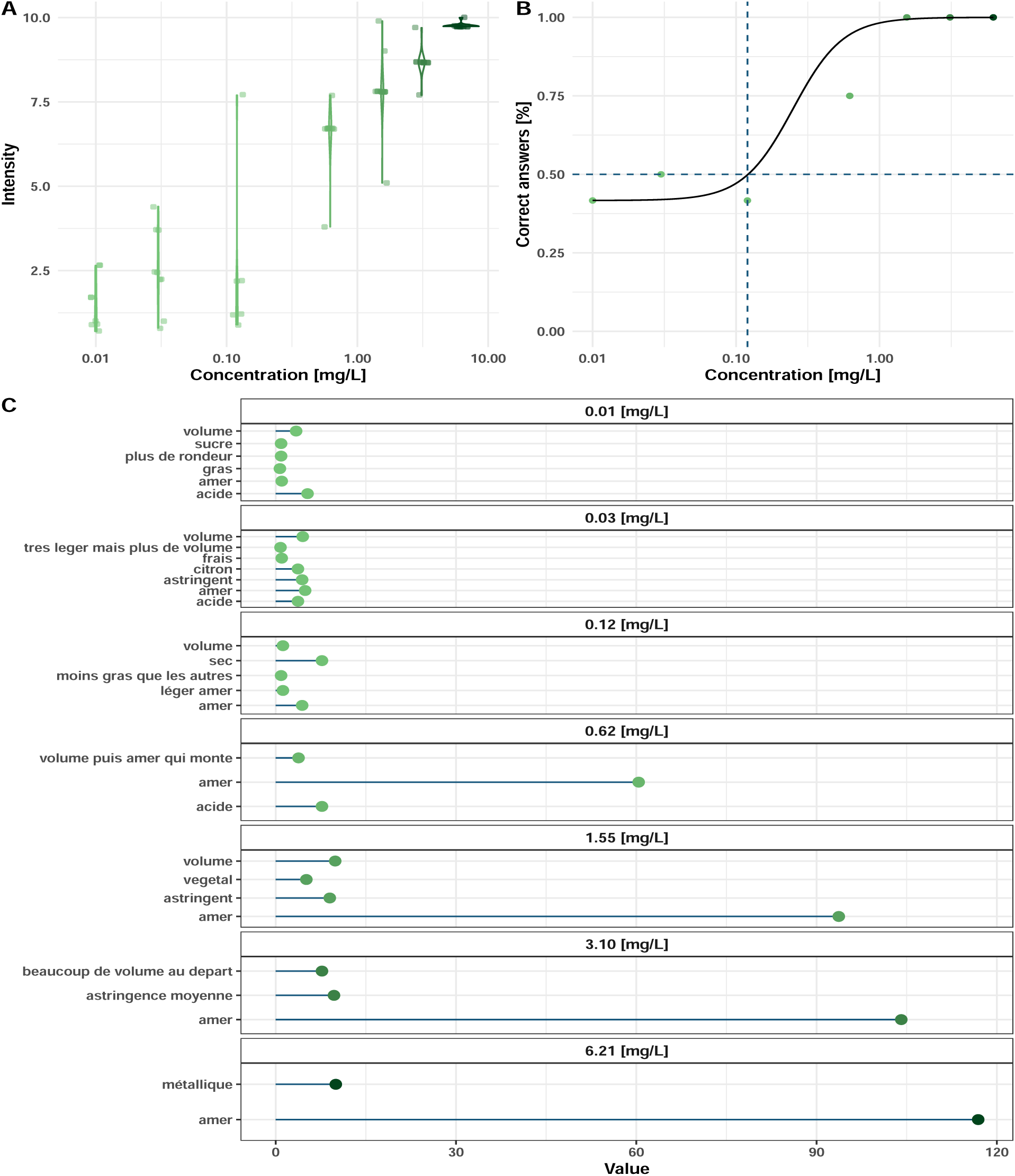
Determination of the concentration used for tasting. Panel A represents the intensity score given to the sample in a triangle test as a function of the enriched extract concentration (only if the answer was correct). Panel B represents the number of correct answers in a triangle test as a function of sample concentration. Finally, panel C represents the occurrence of the descriptors attributed to the sample, multiplied by the intensity given to the sample, as a function of concentration. For more information about triangle test in sensory analysis, see https://www.sensorysociety.org/knowledge/sspwiki/Pages/Triangle%20Test.aspx

**Table S1.** Masses of the MPLC fractions.

| Fraction | Mass [mg] |  | Fraction | Mass [mg] |  | Fraction | Mass [mg] |  | Fraction | Mass [mg] |
| --- | --- | --- | --- | --- | --- | --- | --- | --- | --- | --- |
| M_01 | 0.0 |  | M_22 | 143.0 |  | M_43 | 149.7 |  | M_64 | 51.1 |
| M_02 | 0.0 |  | M_23 | 195.3 |  | M_44 | 127.9 |  | M_65 | 59.8 |
| M_03 | 145.3 |  | M_24 | 199.6 |  | M_45 | 156.7 |  | M_66 | 60.1 |
| M_04 | 91.9 |  | M_25 | 143.8 |  | M_46 | 167.7 |  | M_67 | 69.7 |
| M_05 | 0.0 |  | M_26 | 122.4 |  | M_47 | 167.4 |  | M_68 | 36.1 |
| M_06 | 0.0 |  | M_27 | 115.6 |  | M_48 | 157.8 |  | M_69 | 16.6 |
| M_07 | 0.0 |  | M_28 | 116.2 |  | M_49 | 0.0 |  | M_70 | 33.8 |
| M_08 | 0.0 |  | M_29 | 118.6 |  | M_50 | 150.7 |  | M_71 | 63.3 |
| M_09 | 0.0 |  | M_30 | 115.3 |  | M_51 | 163.0 |  | M_72 | 58.6 |
| M_10 | 0.0 |  | M_31 | 118.4 |  | M_52 | 125.2 |  | M_73 | 146.6 |
| M_11 | 0.0 |  | M_32 | 156.1 |  | M_53 | 126.4 |  | M_74 | 144.4 |
| M_12 | 0.0 |  | M_33 | 184.4 |  | M_54 | 95.0 |  | M_75 | 124.5 |
| M_13 | 4.0 |  | M_34 | 210.1 |  | M_55 | 136.2 |  | M_76 | 111.9 |
| M_14 | 13.4 |  | M_35 | 185.2 |  | M_56 | 152.7 |  | M_77 | 76.2 |
| M_15 | 36.1 |  | M_36 | 141.0 |  | M_57 | 127.1 |  | M_78 | 63.8 |
| M_16 | 26.2 |  | M_37 | 123.9 |  | M_58 | 130.5 |  | M_79 | 60.5 |
| M_17 | 99.1 |  | M_38 | 128.7 |  | M_59 | 115.6 |  | M_80 | 96.8 |
| M_18 | 123.0 |  | M_39 | 144.8 |  | M_60 | 90.7 |  | M_81 | 47.7 |
| M_19 | 64.3 |  | M_40 | 152.7 |  | M_61 | 96.3 |  | M_82 | 15.0 |
| M_20 | 89.6 |  | M_41 | 119.7 |  | M_62 | 38.2 |  |  |  |
| M_21 | 99.1 |  | M_42 | 168.4 |  | M_63 | 61.7 |  | **Total** | **<u>7768.2</u>** |

## References

1. Allard, P.M., Bisson, J., Rutz, A., 2023. Isdb: In silico spectral databases of natural products. URL: https://zenodo.org/record/8287341, doi:10.5281/ZENODO.8287341.

2. Barratt-Fornell, A., Drewnowski, A., 2002. The taste of health: Nature’s bitter gifts. Nutrition Today 37, 144–150. URL: http://dx.doi.org/10.1097/00017285-200207000-00003, doi:10.1097/00017285-200207000-00003.

3. Bass, E., 2023. chromatographR: Chromatographic Data Analysis Toolset. URL: https://ethanbass.github.io/chromatographR/, doi:10.5281/zenodo.6944334. version 0.7.2.

4. Belitz, H., Wieser, H., 1985. Bitter compounds: Occurrence and structure-activity relationships. Food Reviews International 1, 271–354. URL: http://dx.doi.org/10.1080/87559128509540773, doi:10.1080/87559128509540773.

5. Benjamini, Y., Hochberg, Y., 1995. Controlling the false discovery rate: A practical and powerful approach to multiple testing. Journal of the Royal Statistical Society Series B: Statistical Methodology 57, 289–300. URL: http://dx.doi.org/10.1111/j.2517-6161.1995.tb02031.x, doi:10.1111/j.2517-6161.1995.tb02031.x.

6. Choules, M.P., Klein, L.L., Lankin, D.C., McAlpine, J.B., Cho, S.H., Cheng, J., Lee, H., Suh, J.W., Jaki, B.U., Franzblau, S.G., Pauli, G.F., 2018. Residual complexity does impact organic chemistry and drug discovery: The case of rufomyazine and rufomycin. The Journal of Organic Chemistry 83, 6664–6672. URL: http://dx.doi.org/10.1021/acs.joc.8b00988, doi:10.1021/acs.joc.8b00988.

7. Csardi, G., Nepusz, T., 2006. The igraph software package for complex network research. InterJournal Complex Systems, 1695. URL: https://igraph.org.

8. Dührkop, K., Fleischauer, M., Ludwig, M., Aksenov, A.A., Melnik, A.V., Meusel, M., Dorrestein, P.C., Rousu, J., Böcker, S., 2019. Sirius 4: a rapid tool for turning tandem mass spectra into metabolite structure information. Nature Methods 16, 299–302. URL: http://dx.doi.org/10.1038/s41592-019-0344-8, doi:10.1038/s41592-019-0344-8.

9. Dührkop, K., Nothias, L.F., Fleischauer, M., Reher, R., Ludwig, M., Hoffmann, M.A., Petras, D., Gerwick, W.H., Rousu, J., Dorrestein, P.C., Böcker, S., 2020. Systematic classification of unknown metabolites using high-resolution fragmentation mass spectra. Nature Biotechnology 39, 462–471. URL: http://dx.doi.org/10.1038/s41587-020-0740-8, doi:10.1038/s41587-020-0740-8.

10. Dührkop, K., Shen, H., Meusel, M., Rousu, J., Böcker, S., 2015. Searching molecular structure databases with tandem mass spectra using csi:fingerid. Proceedings of the National Academy of Sciences 112, 12580–12585. URL: http://dx.doi.org/10.1073/pnas.1509788112, doi:10.1073/pnas.1509788112.

11. Engel, E., Nicklaus, S., Salles, C., Le Quéré, J.L., 2002. Relevance of omission tests to determine flavour-active compounds in food. Food Quality and Preference 13, 505–513. URL: http://dx.doi.org/10.1016/s0950-3293(02)00136-2, doi:10.1016/s0950-3293(02)00136-2.

12. Fairbanks, M., 2024. tidytable: Tidy Interface to ’data.table’. URL: https://CRAN.R-project.org/package=tidytable. r package version0.11.2.

13. Frerebeau, N., 2024. khroma: Colour Schemes for Scientific Data Visualization. Université Bordeaux Montaigne. Pessac, France. URL: https://packages.tesselle.org/khroma/, doi:10.5281/zenodo.1472077. r package version 1.14.0.

14. Gagolewski, M., 2022. stringi: Fast and portable character string processing in r. Journal of Statistical Software 103. URL: http://dx.doi.org/10.18637/jss.v103.i02, doi:10.18637/jss.v103.i02.

15. Galili, T., 2015. dendextend: an r package for visualizing, adjusting and comparing trees of hierarchical clustering. Bioinformatics 31, 3718–3720. URL: http://dx.doi.org/10.1093/bioinformatics/btv428, doi:10.1093/bioinformatics/btv428.

16. Gatto, L., Gibb, S., Rainer, J., 2020. Msnbase, efficient and elegant r-based processing and visualization of raw mass spectrometry data. Journal of Proteome Research 20, 1063–1069. URL: http://dx.doi.org/10.1021/acs.jproteome.0c00313, doi:10.1021/acs.jproteome. 0c00313.

17. Gaudette, N.J., Pickering, G.J., 2013. Modifying bitterness in functional food systems. Critical Reviews in Food Science and Nutrition 53, 464–481. URL: http://dx.doi.org/10.1080/10408398.2010.542511, doi:10.1080/10408398.2010.542511.

18. Heuckeroth, S., Damiani, T., Smirnov, A., Mokshyna, O., Brungs, C., Korf, A., Smith, J.D., Stincone, P., Dreolin, N., Nothias, L.F., Hyötyläinen, T., Orešič, M., Karst, U., Dorrestein, P.C., Petras, D., Du, X., van der Hooft, J.J.J., Schmid, R., Pluskal, T., 2024. Reproducible mass spectrometry data processing and compound annotation in mzmine 3. Nature Protocols 19, 2597–2641. URL: http://dx.doi.org/10.1038/s41596-024-00996-y, doi:10.1038/s41596-024-00996-y.

19. Hoffmann, M.A., Nothias, L.F., Ludwig, M., Fleischauer, M., Gentry, E.C., Witting, M., Dorrestein, P.C., Dührkop, K., Böcker, S., 2021. High-confidence structural annotation of metabolites absent from spectral libraries. Nature Biotechnology 40, 411–421. URL: http://dx.doi.org/10.1038/s41587-021-01045-9, doi:10.1038/s41587-021-01045-9.

20. Hollander, M., A. Wolfe, D., Chicken, E., 2015. Nonparametric Statistical Methods. Wiley. URL: http://dx.doi.org/10.1002/9781119196037, doi:10.1002/9781119196037.

21. Hostettmann-Kaldas, M., Hostettmann, K., Sticher, O., 1981. Xanthones, flavones and secoiridoids of american gentiana species. Phytochemistry 20, 443–446. URL: http://dx.doi.org/10.1016/s0031-9422(00)84162-x, doi:10.1016/s0031-9422(00)84162-x.

22. Hulstaert, N., Shofstahl, J., Sachsenberg, T., Walzer, M., Barsnes, H., Martens, L., Perez-Riverol, Y., 2019. Thermorawfileparser: Modular, scalable, and cross-platform raw file conversion. Journal of Proteome Research 19, 537–542. URL: http://dx.doi.org/10.1021/acs.jproteome.9b00328, doi:10.1021/acs.jproteome.9b00328.

23. Husson, F., Le, S., Cadoret, M., 2023. SensoMineR: Sensory Data Analysis. URL: https://CRAN.R-project.org/package=SensoMineR. r package version 1.27.

24. Husson, F., Pagès, J., 2006. Indscal model: geometrical interpretation and methodology. Computational Statistics & Data Analysis 50, 358–378. URL: http://dx.doi.org/10.1016/j.csda.2004.08.005, doi:10.1016/j.csda.2004.08.005.

25. Jarmusch, A.K., Wang, M., Aceves, C.M., Advani, R.S., Aguirre, S., Aksenov, A.A., Aleti, G., Aron, A.T., Bauermeister, A., Bolleddu, S., Bouslimani, A., Caraballo Rodriguez, A.M., Chaar, R., Coras, R., Elijah, E.O., Ernst, M., Gauglitz, J.M., Gentry, E.C., Husband, M., Jarmusch, S.A., Jones, K.L., Kamenik, Z., Le Gouellec, A., Lu, A., McCall, L.I., McPhail, K.L., Meehan, M.J., Melnik, A.V., Menezes, R.C., Montoya Giraldo, Y.A., Nguyen, N.H., Nothias, L.F., Nothias-Esposito, M., Panitchpakdi, M., Petras, D., Quinn, R.A., Sikora, N., van der Hooft, J.J.J., Vargas, F., Vrbanac, A., Weldon, K.C., Knight, R., Bandeira, N., Dorrestein, P.C., 2020. Redu: a framework to find and reanalyze public mass spectrometry data. Nature Methods 17, 901–904. URL: http://dx.doi.org/10.1038/s41592-020-0916-7, doi:10.1038/s41592-020-0916-7.

26. Jiang, A., 2024. NMRphasing: Phase Error Correction and Baseline Correction for One Dimensional (’1D’) ’NMR’ Data. URL: https://CRAN.R-project.org/package=NMRphasing. r package version 1.0.5.

27. Joshi, K., Hankey, A., Patwardhan, B., 2006. Traditional phytochemistry: Identification of drug by ‘taste’. Evidence-Based Complementary and Alternative Medicine 4, 145–148. URL: http://dx.doi.org/10.1093/ecam/nel064, doi:10.1093/ecam/nel064.

28. Kassambara, A., 2023. ggpubr: ’ggplot2’ Based Publication Ready Plots. URL: https://CRAN.R-project.org/package=ggpubr. r package version 0.6.0.

29. Kawahara, N., Masuda, K., Sekita, S., Satake, M., 2001. A new secoiridoid glucoside, amaronitidin, from the peruvian folk medicine “hercampuri” (gentianella nitida). Chemical and Pharmaceutical Bulletin 49, 771–772. URL: http://dx.doi.org/10.1248/cpb.49.771, doi:10.1248/cpb.49.771.

30. Kumar, V., Van Staden, J., 2016. A review of swertia chirayita (gentianaceae) as a traditional medicinal plant. Frontiers in Pharmacology 6. URL: http://dx.doi.org/10.3389/fphar.2015.00308, doi:10.3389/fphar.2015.00308.

31. Lance, G.N., Williams, W.T., 1966. Computer programs for hierarchical polythetic classification (“similarity analyses”). The Computer Journal 9, 60–64. URL: http://dx.doi.org/10.1093/comjnl/9.1.60, doi:10.1093/comjnl/9.1.60.

32. Lê, S., Josse, J., Husson, F., 2008. Factominer: Anrpackage for multivariate analysis. Journal of Statistical Software 25. URL: http://dx.doi.org/10.18637/jss.v025.i01, doi:10.18637/jss.v025.i01.

33. Ludwig, M., Nothias, L.F., Dührkop, K., Koester, I., Fleischauer, M., Hoffmann, M.A., Petras, D., Vargas, F., Morsy, M., Aluwihare, L., Dorrestein, P.C., Böcker, S., 2020. Database-independent molecular formula annotation using gibbs sampling through zodiac. Nature Machine Intelligence 2, 629–641. URL: http://dx.doi.org/10.1038/s42256-020-00234-6, doi:10.1038/s42256-020-00234-6.

34. Madrid-Gambin, F., Oller-Moreno, S., Fernandez, L., Bartova, S., Giner, M.P., Joyce, C., Ferraro, F., Montoliu, I., Moco, S., Marco, S., 2020. Alpsnmr: an r package for signal processing of fully untargeted nmr-based metabolomics. Bioinformatics 36, 2943–2945. URL: http://dx.doi.org/10.1093/bioinformatics/btaa022, doi:10.1093/bioinformatics/btaa022.

35. Martens, L., Chambers, M., Sturm, M., Kessner, D., Levander, F., Shofstahl, J., Tang, W.H., Römpp, A., Neumann, S., Pizarro, A.D., Montecchi-Palazzi, L., Tasman, N., Coleman, M., Reisinger, F., Souda, P., Hermjakob, H., Binz, P.A., Deutsch, E.W., 2011. mzml—a community standard for mass spectrometry data. Molecular & Cellular Proteomics 10, R110.000133. URL: http://dx.doi.org/10.1074/mcp.R110.000133, doi:10.1074/mcp.r110.000133.

36. McAlpine, J.B., Chen, S.N., Kutateladze, A., MacMillan, J.B., Appendino, G., Barison, A., Beniddir, M.A., Biavatti, M.W., Bluml, S., Boufridi, A., Butler, M.S., Capon, R.J., Choi, Y.H., Coppage, D., Crews, P., Crimmins, M.T., Csete, M., Dewapriya, P., Egan, J.M., Garson, M.J., Genta-Jouve, G., Gerwick, W.H., Gross, H., Harper, M.K., Hermanto, P., Hook, J.M., Hunter, L., Jeannerat, D., Ji, N.Y., Johnson, T.A., Kingston, D.G.I., Koshino, H., Lee, H.W., Lewin, G., Li, J., Linington, R.G., Liu, M., McPhail, K.L., Molinski, T.F., Moore, B.S., Nam, J.W., Neupane, R.P., Niemitz, M., Nuzillard, J.M., Oberlies, N.H., Ocampos, F.M.M., Pan, G., Quinn, R.J., Reddy, D.S., Renault, J.H., Rivera-Chávez, J., Robien, W., Saunders, C.M., Schmidt, T.J., Seger, C., Shen, B., Steinbeck, C., Stuppner, H., Sturm, S., Taglialatela-Scafati, O., Tantillo, D.J., Verpoorte, R., Wang, B.G., Williams, C.M., Williams, P.G., Wist, J., Yue, J.M., Zhang, C., Xu, Z., Simmler, C., Lankin, D.C., Bisson, J., Pauli, G.F., 2019. The value of universally available raw nmr data for transparency, reproducibility, and integrity in natural product research. Natural Product Reports 36, 35–107. URL: http://dx.doi.org/10.1039/c7np00064b, doi:10.1039/c7np00064b.

37. Müller, K., Wickham, H., 2023. tibble: Simple Data Frames. URL: https://CRAN.R-project.org/package=tibble. r package version 3.2.1.

38. Pagès, J., 2003. Recueil direct de distances sensorielles: application à l’évaluation de dix vins blancs du val-de-loire. Sciences des Aliments 23, 679–688. URL: http://dx.doi.org/10.3166/sda.23.679-688, doi:10.3166/sda.23.679-688.

39. Pagès, J., 2005. Collection and analysis of perceived product inter-distances using multiple factor analysis: Application to the study of 10 white wines from the loire valley. Food Quality and Preference 16, 642–649. URL: http://dx.doi.org/10.1016/j.foodqual.2005.01.006, doi:10.1016/j.foodqual.2005.01.006.

40. Pedersen, T.L., 2024. ggraph: An Implementation of Grammar of Graphics for Graphs and Networks. URL: https://CRAN.R-project.org/package=ggraph. r package version 2.2.1.

41. Phoboo, S., Pinto, M.D.S., Barbosa, A.C.L., Sarkar, D., Bhowmik, P.C., Jha, P.K., Shetty, K., 2012. Phenolic-linked biochemical rationale for the anti-diabetic properties of swertia chirayita (roxb. ex flem.) karst. Phytotherapy Research 27, 227–235. URL: http://dx.doi.org/10.1002/ptr.4714, doi:10.1002/ptr.4714.

42. Pluskal, T., Castillo, S., Villar-Briones, A., Orešič, M., 2010. Mzmine 2: Modular framework for processing, visualizing, and analyzing mass spectrometry-based molecular profile data. BMC Bioinformatics 11. URL: http://dx.doi.org/10.1186/1471-2105-11-395, doi:10.1186/1471-2105-11-395.

43. Rainer, J., Vicini, A., Salzer, L., Stanstrup, J., Badia, J.M., Neumann, S., Stravs, M.A., Verri Hernandes, V., Gatto, L., Gibb, S., Witting, M., 2022. A modular and expandable ecosystem for metabolomics data annotation in r. Metabolites 12, 173. URL: http://dx.doi.org/10.3390/metabo12020173, doi:10.3390/metabo12020173.

44. Rojas, C., Ballabio, D., Pacheco Sarmiento, K., Pacheco Jaramillo, E., Mendoza, M., García, F., 2022. Chemtastesdb: A curated database of molecular tastants. Food Chemistry: Molecular Sciences 4, 100090. URL: http://dx.doi.org/10.1016/j.fochms.2022.100090, doi:10.1016/j.fochms.2022.100090.

45. Rotzoll, N., Dunkel, A., Hofmann, T., 2006. Quantitative studies, taste reconstitution, and omission experiments on the key taste compounds in morel mushrooms (morchella deliciosa fr.). Journal of Agricultural and Food Chemistry 54, 2705–2711. URL: http://dx.doi.org/10.1021/jf053131y, doi:10.1021/jf053131y.

46. Royston, J.P., 1982. An extension of shapiro and wilk’s w test for normality to large samples. Applied Statistics 31, 115. URL: http://dx.doi.org/10.2307/2347973, doi:10.2307/2347973.

47. Rutz, A., 2024a. cascade: Contextualizing untargeted annotation with semi-quantitative charged aerosol detection for pertinent characterization of natural extracts. URL: https://zenodo.org/doi/10.5281/zenodo.14515159, doi:10.5281/ZENODO.14515159.

48. Rutz, A., 2024b. tima: Taxonomically informed metabolite annotation. URL: https://zenodo.org/doi/10.5281/zenodo.14515116, doi:10.5281/ZENODO.14515116.

49. Rutz, A., 2025. sapid: A strategy to analyze plant extracts taste in depth. URL: https://zenodo.org/doi/10.5281/zenodo.14616395, doi:10.5281/ZENODO.14616395.

50. Rutz, A., Bisson, J., Allard, P.M., 2023. The lotus initiative for open natural products research: frozen dataset union wikidata (with metadata). URL: https://zenodo.org/record/7534071, doi:10.5281/ZENODO.7534071.

51. Rutz, A., Dounoue-Kubo, M., Ollivier, S., Bisson, J., Bagheri, M., Saesong, T., Ebrahimi, S.N., Ingkaninan, K., Wolfender, J.L., Allard, P.M., 2019. Taxonomically informed scoring enhances confidence in natural products annotation. Frontiers in Plant Science 10. URL: http://dx.doi.org/10.3389/fpls.2019.01329, doi:10.3389/fpls.2019.01329.

52. Rutz, A., Marcourt, L., Wolfender, J.L., 2024. Nmr data of ethanolic extract and fractions of swertia chirayita. URL: https://zenodo.org/doi/10.5281/zenodo.14414271, doi:10.5281/ZENODO.14414271.

53. Rutz, A., Wolfender, J.L., 2023. Automated composition assessment of natural extracts: Untargeted mass spectrometry-based metabolite profiling integrating semiquantitative detection. Journal of Agricultural and Food Chemistry 71, 18010–18023. URL: http://dx.doi.org/10.1021/acs.jafc.3c03099, doi:10.1021/acs.jafc.3c03099.

54. Saha, P., Das, S., 2010. Highlighting the anti-carcinogenic potential of an ayurvedic medicinal plant, Swertia Chirata. Asian Pacific journal of cancer prevention : APJCP 11, 1445–9.

55. Schmid, R., Petras, D., Nothias, L.F., Wang, M., Aron, A.T., Jagels, A., Tsugawa, H., Rainer, J., Garcia-Aloy, M., Dührkop, K., Korf, A., Pluskal, T., Kameník, Z., Jarmusch, A.K., Caraballo-Rodríguez, A.M., Weldon, K.C., Nothias-Esposito, M., Aksenov, A.A., Bauermeister, A., Albarracin Orio, A., Grundmann, C.O., Vargas, F., Koester, I., Gauglitz, J.M., Gentry, E.C., Hövelmann, Y., Kalinina, S.A., Pendergraft, M.A., Panitchpakdi, M., Tehan, R., Le Gouellec, A., Aleti, G., Mannochio Russo, H., Arndt, B., Hübner, F., Hayen, H., Zhi, H., Raffatellu, M., Prather, K.A., Aluwihare, L.I., Böcker, S., McPhail, K.L., Humpf, H.U., Karst, U., Dorrestein, P.C., 2021. Ion identity molecular networking for mass spectrometry-based metabolomics in the gnps environment. Nature Communications 12. URL: http://dx.doi.org/10.1038/s41467-021-23953-9, doi:10.1038/s41467-021-23953-9.

56. Sharma, Y.P., Bhardwaj, C., Sharma, R., Thakur, P., Sharma, R., 2022. A new rp-hplc method for simultaneous determination of amaroswerin, amarogentin and andrographolide in a herbal drug “chirayata”. Journal of Chromatographic Science 61, 172–176. URL: http://dx.doi.org/10.1093/chromsci/bmac018, doi:10.1093/chromsci/bmac018.

57. Shukla, S., Bafna, K., Sundar, D., Thorat, S.S., 2014. The bitter barricading of prostaglandin biosynthesis pathway: Understanding the molecular mechanism of selective cyclooxygenase-2 inhibition by amarogentin, a secoiridoid glycoside from swertia chirayita. PLoS ONE 9, e90637. URL: http://dx.doi.org/10.1371/journal.pone.0090637, doi:10.1371/journal.pone.0090637.

58. Sjoberg, D., 2020. ggbump: Bump Chart and Sigmoid Curves. URL: https://CRAN.R-project.org/package=ggbump. r package version 0.1.0.

59. Stravs, M.A., Dührkop, K., Böcker, S., Zamboni, N., 2022. Msnovelist: de novo structure generation from mass spectra. Nature Methods 19, 865–870. URL: http://dx.doi.org/10.1038/s41592-022-01486-3, doi:10.1038/s41592-022-01486-3.

60. Suryawanshi, S., Mehrotra, N., Asthana, R.K., Gupta, R.C., 2006. Liquid chromatography/tandem mass spectrometric study and analysis of xanthone and secoiridoid glycoside composition of swertia chirata, a potent antidiabetic. Rapid Communications in Mass Spectrometry 20, 3761–3768. URL: http://dx.doi.org/10.1002/rcm.2795, doi:10.1002/rcm.2795.

61. Vaughan, D., Dancho, M., 2022. furrr: Apply Mapping Functions in Parallel using Futures. URL: https://CRAN.R-project.org/package=furrr. r package version 0.3.1.

62. Wang, F., Liigand, J., Tian, S., Arndt, D., Greiner, R., Wishart, D.S., 2021. Cfm-id 4.0: More accurate esi-ms/ms spectral prediction and compound identification. Analytical Chemistry 93, 11692–11700. URL: http://dx.doi.org/10.1021/acs.analchem.1c01465, doi:10.1021/acs.analchem.1c01465.

63. Ward, J.H., 1963. Hierarchical grouping to optimize an objective function. Journal of the American Statistical Association 58, 236–244. URL:http://dx.doi.org/10.1080/01621459.1963.10500845, doi:10.1080/01621459.1963.10500845.

64. Wickham, H., 2016. ggplot2: Elegant Graphics for Data Analysis. Springer-Verlag New York. URL: https://ggplot2.tidyverse.org.

65. Wickham, H., 2023. forcats: Tools for Working with Categorical Variables (Factors). URL: https://CRAN.R-project.org/package=forcats. r package version 1.0.0.

66. Wickham, H., Bryan, J., 2023. readxl: Read Excel Files. URL: https://CRAN.R-project.org/package=readxl. r package version 1.4.3.

67. Wickham, H., Pedersen, T.L., Seidel, D., 2023. scales: Scale Functions for Visualization. URL: https://CRAN.R-project.org/package=scales. r package version 1.3.0.

68. Wilkins, D., 2023. treemapify: Draw Treemaps in ’ggplot2’. URL: https://CRAN.R-project.org/package=treemapify. r package version 2.5.6.

69. Wolfender, J.L., Hamburger, M., Hostettmann, K., Msonthi, J.D., Mavi, S., 1993. Search for bitter principles in chironia species by lc-ms and isolation of a new secoiridoid diglycoside from chironia krebsii. Journal of Natural Products 56, 682–689. URL: http://dx.doi.org/10.1021/np50095a004, doi:10.1021/np50095a004.

70. Wölfle, U., Schempp, C.M., 2018. Bitterstoffe–von der traditionellen verwendung bis zum einsatz an der haut. Zeitschrift für Phytotherapie 39, 210–215. URL: http://dx.doi.org/10.1055/a-0654-1711, doi:10.1055/a-0654-1711.

71. Xing, S., Shen, S., Xu, B., Li, X., Huan, T., 2023. Buddy: molecular formula discovery via bottom-up ms/ms interrogation. Nature Methods 20, 881–890. URL: http://dx.doi.org/10.1038/s41592-023-01850-x, doi:10.1038/s41592-023-01850-x.

72. Yan, J., Tong, H., 2022. An overview of bitter compounds in foodstuffs: Classifications, evaluation methods for sensory contribution, separation and identification techniques, and mechanism of bitter taste transduction. Comprehensive Reviews in Food Science and Food Safety 22, 187–232. URL: http://dx.doi.org/10.1111/1541-4337.13067, doi:10.1111/1541-4337.13067.

73. Zhao, A., Jeffery, E.H., Miller, M.J., 2022. Is bitterness only a taste? the expanding area of health benefits of brassica vegetables and potential for bitter taste receptors to support health benefits. Nutrients 14, 1434. URL: http://dx.doi.org/10.3390/nu14071434, doi:10.3390/nu14071434.

